# The termination of UHRF1-dependent PAF15 ubiquitin signaling is regulated by USP7 and ATAD5

**DOI:** 10.1101/2022.05.30.494002

**Authors:** Ryota Miyashita, Atsuya Nishiyama, Yoshie Chiba, Satomi Kori, Norie Kato, Chieko Konishi, Soichiro Kumamoto, Hiroko Kozuka-Hata, Masaaki Oyama, Yoshitaka Kawasoe, Toshiki Tsurimoto, Tatsuro S Takahashi, Kyohei Arita, Makoto Nakanishi

**Affiliations:** Division of Cancer Cell Biology, The Institute of Medical Science, The University of Tokyo, 4-6-1 Shirokanedai, Tokyo, Japan; Medical Proteomics Laboratory, The Institute of Medical Science, The University of Tokyo, 4-6-1 Shirokanedai, Tokyo, Japan; Structural Biology Laboratory, Graduate School of Medical Life Science, Yokohama City University, 1-7-29, Suehiro-cho, Kanagawa, Japan; Laboratory of Chromosome Biology, Department of Biology, Faculty of Science, Kyushu University, 744, Motooka, Nishi-ku, Fukuoka, Japan

## Abstract

UHRF1-dependent ubiquitin signaling plays an integral role in the regulation of maintenance DNA methylation. UHRF1 catalyzes transient dual mono-ubiquitylation of PAF15 (PAF15Ub2), which regulates the localization and activation of DNMT1 at DNA methylation sites during DNA replication. Although the initiation of UHRF1-mediated PAF15 ubiquitin signaling has been relatively well characterized, mechanisms underlying its termination and how they are coordinated with the completion of maintenance DNA methylation have not yet been clarified. This study shows that deubiquitylation by USP7 and unloading by ATAD5 (ELG1 in yeast) are pivotal processes for the removal of PAF15 from chromatin. On replicating chromatin, USP7 specifically interacts with PAF15Ub2 in a complex with DNMT1. USP7 depletion or inhibition of the interaction between USP7 and PAF15 results in abnormal accumulation of PAF15Ub2 on chromatin. Furthermore, we also find that the non-ubiquitylated form of PAF15 (PAF15Ub0) is removed from chromatin in an ATAD5-dependent manner. PAF15Ub2 was retained at high levels on chromatin when the catalytic activity of DNMT1 was inhibited, suggesting that the completion of maintenance DNA methylation is essential for termination of UHRF1-mediated ubiquitin signaling. This finding provides a molecular understanding of how the maintenance DNA methylation machinery is disassembled at the end of the S phase.

## Introduction

DNA methylation at CpG dinucleotide is an epigenetic modification that regulates various biological processes, including gene silencing, genome stability, cellular development, and differentiation (Greenberg & Bourc’his, 2019; Jones, 2012; Moore et al., 2013). DNA methylation is stably maintained during cell proliferation (Jones & Liang, 2009; Petryk et al., 2021). DNA methyltransferase 1 (DNMT1) plays a key role in the maintenance of DNA methylation by catalyzing the conversion of hemi-methylated DNA to a fully methylated state (Edwards et al., 2017). In addition, recent studies have also suggested the potential de novo function of DNMT1 (Haggerty et al., 2021; Li, Jialun et al., 2020; Li, Yingfeng et al., 2018). Besides the C-terminal catalytic domain, DNMT1 contains several regulatory regions, including Proliferating cell nuclear antigen (PCNA)-interacting protein motif (PIP-box), replication foci targeting sequence (RFTS), a CXXC zinc finger domain, and two bromo-adjacent homology (BAH) domains (Lyko, 2018). DNMT1 specifically localizes at DNA methylation sites dependently on the RFTS domain, which interacts with dual mono-ubiquitylated histone H3 (H3Ub2) or PAF15 (PAF15Ub2) (Ishiyama et al., 2017; Nishiyama et al., 2020; Qin et al., 2015). The RFTS domain of DNMT1 also shows preferential H3K9me3 binding over H3K9me0 to enhance the interaction with H3Ub2 (Ren et al, 2020). These interactions also cause release of autoinhibition and enzymatic activation of DNMT1, presumably via the conformational change (Ishiyama et al., 2017; Mishima et al., 2020; Syeda et al., 2011; Takeshita et al., 2011; Zhang, Liu, et al., 2015).

Dual monoubiquitylation of histone H3 and PAF15 is catalyzed by an E3 ubiquitin ligase Ubiquitin-like containing PHD and RING finger domains 1 (UHRF1), also known as NP95 or ICBP90 (Karg et al., 2017; Nishiyama et al., 2013). UHRF1 binds specifically to hemi-methylated DNA via its SET and RING-associated (SRA) domain (Arita et al., 2008; Avvakumov et al., 2008; Hashimoto et al., 2008) and plays an essential role for the DNMT1 recruitment to sites of DNA methylation (Bostick et al., 2007; Sharif et al., 2007). The E3 ubiquitin ligase activity of UHRF1 is enhanced by binding to hemi-methylated DNA (Harrison et al., 2016), and mutations in the RING finger domain, which is responsible for ubiquitin ligase activity, impair the localization of DNMT1 to methylation sites and maintenance DNA methylation (Nishiyama et al., 2013; Qin et al., 2015). The N-terminal Ubiquitin-like (UBL) domain promotes interaction with the E2 enzyme, Ubch5/UBE2D (DaRosa et al., 2018; Foster et al., 2018). The plant homeodomain (PHD) and tandem Tudor (TTD) domains are responsible for recognizing and binding to the N-terminal portion of histone H3 and PAF15 (Arita et al., 2012; Nishiyama et al., 2020; Rajakumara et al., 2011; Rothbart et al., 2012). UHRF1 dissociates from chromatin upon conversion of hemi-methylated DNA to fully methylated DNA, leading to the inactivation of UHRF1-dependent ubiquitin signaling (Nishiyama et al., 2020).

PAF15 is a PCNA-binding protein (De Biasio et al., 2015; Emanuele et al., 2011; Karg et al., 2017; Nishiyama et al., 2020; Yu et al., 2001) and transiently binds to chromatin during S phase in PCNA- and DNA replication-dependent manner (Nishiyama et al., 2020). In addition, inhibition of UHRF1-dependent PAF15 ubiquitylation significantly impairs PAF15 chromatin binding (Karg et al., 2017; Nishiyama et al., 2020), suggesting that ubiquitylation of PAF15 plays an important role not only in its interaction with DNMT1 but also in its own chromatin binding. Given that more than 80% of CpG methylation on the genome is maintained by DNA replication-coupled maintenance (Charlton et al., 2018; Ming et al., 2020) and that PAF15Ub2 is a key regulator of replication-coupled DNMT1 chromatin recruitment, a regulatory mechanism for PAF15 ubiquitylation is critical for faithful propagation of DNA methylation patterns. Inefficient termination of PAF15 ubiquitin signaling will result in overloading of DNMT1 and unregulated DNA methylation, which is frequently observed in various types of tumors. However, it is not fully understood how the termination of PAF15 ubiquitin signaling is regulated during the process of maintenance DNA methylation.

Protein ubiquitylation is a reversible post-translational modification (Komander & Rape, 2012). Among nearly 100 deubiquitylating enzymes, USP7 (Ubiquitin-Specific Protease 7, also known as HAUSP) has been shown to accumulate at DNA methylation sites in a complex with DNMT1 or UHRF1 (Felle et al., 2011; Ma et al., 2012; Qin et al., 2011; Yamaguchi et al., 2017; Zhang, Rothbart, et al., 2015). While it has been reported that USP7 promotes efficient maintenance of DNA methylation through stabilization of DNMT1 and UHRF1 by preventing their polyubiquitylation and proteasomal degradation (Cheng, Yang, et al., 2015; Du et al., 2010), recent studies have shown that USP7 also modulates the level of ubiquitinated histone H3 and histone H2B on chromatin (Li et al., 2020; Yamaguchi et al., 2017). However, it remains unclear whether USP7 also regulates the PAF15 ubiquitylation.

In this report, we set out to study the molecular mechanism of PAF15 chromatin unloading to understand how the termination of maintenance DNA methylation is regulated. Using the cell-free system derived from *Xenopus* egg extracts that recapitulate the processes of maintenance DNA methylation, we demonstrate that the unloading of PAF15Ub2 is regulated by the two regulatory mechanisms, namely USP7-dependent deubiquitylation and unloading of PAF15 by ATPase family AAA domain-containing protein 5 (ATAD5). We also find that PAF15 unloading is tightly coordinated with the completion of maintenance DNA methylation and requires the release of UHRF1 from chromatin. Finally, co-depletion of USP7 and ATAD5 from egg extracts results in an elevated global DNA methylation. We propose that timely inactivation of PAF15 is critical for the faithful inheritance of DNA methylation patterns.

## Results

### Identification of USP7 as a PAF15 binding protein

UHRF1-dependent ubiquitin signaling plays a critical role in PAF15 chromatin binding. To test whether the deubiquitylation (DUB) activity is required for termination of PAF15 ubiquitylation signaling, we employed Ubiquitin vinyl sulfone (UbVS), a pan DUB inhibitor (Borodovsky et al., 2001). In *Xenopus* egg extracts, sperm chromatin added to egg cytoplasm assembles a functional nucleus and undergoes chromosomal replication (Blow & Laskey, 2016). As it has been reported that treatment of *Xenopus* egg extracts with UbVS inhibits ubiquitin turnover, resulting in depletion of free ubiquitin (Dimova et al., 2012), we also added excess free ubiquitin before incubation with sperm chromatin to activate ubiquitylation pathways. As previously reported, PAF15 underwent dual mono-ubiquitylation on chromatin during S phase and then dissociated from chromatin (Nishiyama et al., 2020; Povlsen et al., 2012). The addition of UbVS alone significantly delayed PAF15 and DNMT1 chromatin loading, confirming the importance of ubiquitin signaling for the initiation of maintenance of DNA methylation (Fig. 1A, S1A). In contrast, inhibition of DUB by UbVS plus excess free ubiquitin led to enhanced and prolonged chromatin association of DNMT1 and PAF15, indicating that the DUB activity was required for the termination of maintenance DNA methylation in egg extracts.

**Figure 1.**
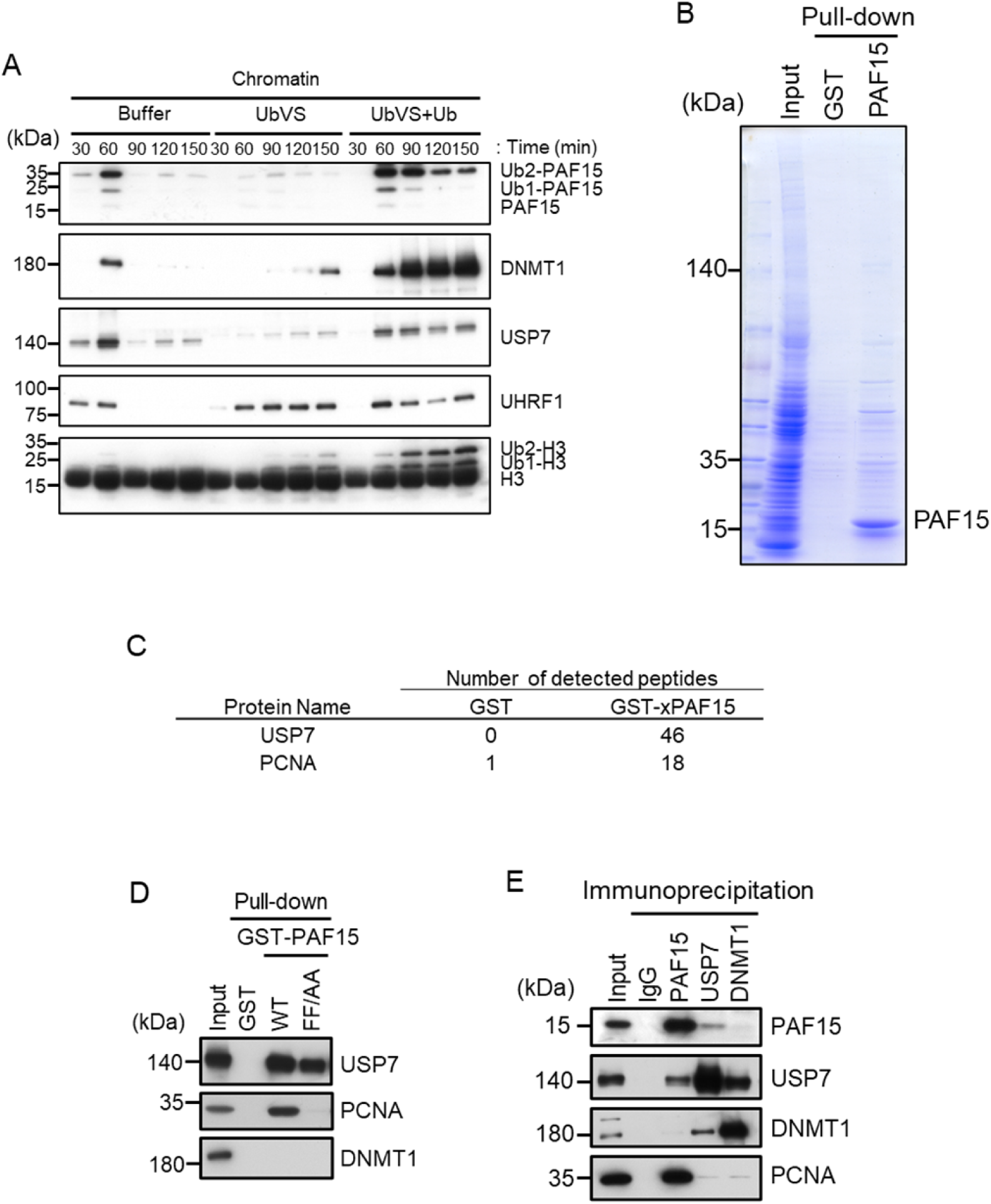
USP7 was identified as a PAF15 binding protein. (A) Sperm chromatin was added to interphase egg extracts supplemented with either buffer (+Buffer), 20 µM UbVS (+UbVS), or 20 µM UbVS and 58 µM ubiquitin (+UbVS+Ub). Samples were analyzed by immunoblotting using the antibodies indicated. (B) Proteins pull-downed from interphase egg extracts by GST and GST-PAF15 were stained by CBB. (C) The samples from GST-PAF15 pulldown were analyzed by nanoLC-MS/MS. Selected proteins were indicated in the table. (D) GST pull-down assay was performed by GST or GST-PAF15 wild-type (WT) or PIP mutant (FF/AA), and the samples were analyzed by immunoblotting using the antibodies indicated. (E) Immunoprecipitation was performed by PAF15, USP7 and DNMT1 antibodies-bound beads, and the samples were analyzed by immunoblotting using the antibodies indicated. Source Data are provided as Figure 1-source data.

To examine the possibility that PAF15 interacts proteins related to deubiquitylation, we performed glutathione-S-transferase (GST)-PAF15 pull-down and nanoflow liquid chromatography-tandem mass spectrometry (nanoLC-MS/MS) for identification of proteins. Sepharose beads bound to GST or GST-PAF15 were incubated with *Xenopus* interphase egg extracts. Beads-bound proteins were eluted by cleavage of GST with thrombin protease (Fig. 1B). Recovered proteins were identified by nanoLC-MS/MS (Fig. 1C, Supplementary Tables 1 and 2). This analysis confirmed the PAF15 binding with PCNA and revealed the interaction between PAF15 and USP7. The interaction between GST-PAF15 and USP7 was also validated by immunoblotting using USP7 specific antibodies. Mutations of two phenylalanine to alanine within the consensus PIP sequence of PAF15 abolished the interaction with PCNA, but not USP7, suggesting that the PAF15-USP7 interaction is independent of PCNA (Fig. 1D, S1B). This interaction was further demonstrated by reciprocal immunoprecipitation and western blotting experiments for endogenous proteins in egg extracts using anti-PAF15 and USP7 antibodies (Fig. 1E). Our results indicate that USP7 has an activity to interact with PAF15 although it was still unclear at this stage whether other protein(s) might be involved in this interaction.

### PAF15 associates with USP7 through the TRAF and UBL2 domains

It has been reported that the binding of USP7 to substrate proteins involves two distinct domains: one is the N-terminal TRAF (TNF-receptor-associated factors-like) domain and the other is the C-terminal UBL (Ubiquitin-like) domain (Al-Eidan et al., 2020). Previous reports have shown that the recognition of the P/A/ExxS motif via the binding pocket within the TRAF domain of USP7 is important for the interaction with many substrate proteins such as p53, MDM2, and MCM-BP (Hu et al., 2006; Jagannathan et al., 2014; Sheng et al., 2006). Meanwhile, it has also been reported that the UBL2 domain recognizes KxxxK motifs that interact with acidic surface patches within the UBL2 (Cheng, Li, et al., 2015). We searched for these motifs in PAF15 and found two P/AxxS motifs (^76^PSTS^79^, ^94^AGGS^97^) and one KxxxK motif (^101^KKPRK^105^) (Fig. 2A). We then tested whether these sequences serve as binding sites for USP7 by mutating serine residues in the P/AxxS motif and two lysine residues in the KxxxK motif to alanine (rPAF15 SA and KA, respectively), and combining these mutations to produce a triple mutant (rPAF15 SAKA). As described above, GST-PAF15 was able to pull down both USP7 and PCNA from *Xenopus* interphase egg extracts (Fig. 2B). The PIP-box mutant of PAF15 lost interaction with PCNA but retained binding to USP7. In contrast, mutations in the P/AxxS or KxxxK motifs reduced the binding of USP7 to GST-PAF15. Furthermore, the triple mutations completely lost the binding of PAF15 to USP7. These results suggest that USP7 interacts with PAF15 through both the TRAF domain and the UBL2 domain.

**Figure 2.**
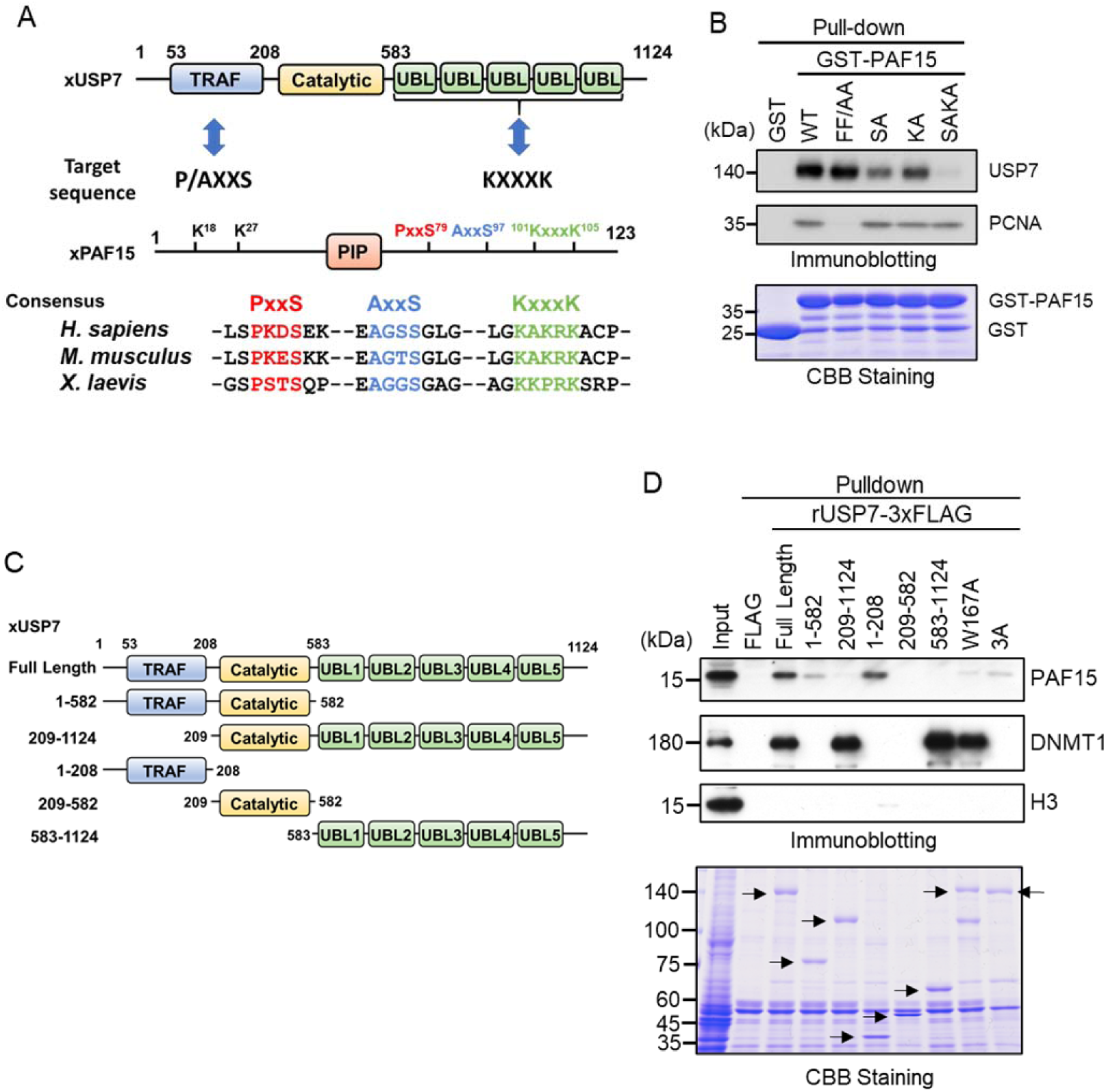
PAF15 associates with the TRAF and UBL1-2 domains of USP7. **(A)** Schematic illustration of PAF15-USP7 binding experiment. USP7 recognizes P/AxxS or KxxxK motifs in its substrates via TRAF or UBL domains, respectively. PAF15 has three motifs, and alanine mutations were introduced at S79, S97, K101 and K105. **(B)** GST pull-down from the interphase egg extracts using GST-PAF15, P/AxxS mutant (S79A/S97A; SA), KxxxK mutant (K101A/K105A; KA), and triple mutant (SAKA). The samples were analyzed by immunoblotting using the antibodies indicated. Purified GST or GST-PAF15 mutants used in pull-down assay were stained using CBB. **(C)** Schematic illustration of rUSP7 truncation mutants employed in (D). **(D)** FLAG pull-down from interphase egg extracts using rUSP7-3xFLAG mutants presented in (C), W167A and 3A point mutants. USP7 3A: D780A/E781A/D786A. The samples were analyzed by immunoblotting using the antibodies indicated. Samples were also stained by CBB. Arrowheads indicate rUSP7 truncation mutants and point mutants. Source Data are provided as Figure 2-source data.

Next, we confirmed the requirement of the TRAF and UBL2 domains of USP7 for the interaction with PAF15 by performing pull-down experiments from egg extracts using 3xFLAG-tagged-rUSP7 and its mutants expressed in insect cells (Fig. 2C). The TRAF domain (residues 1-208) interacted efficiently with PAF15, while the catalytic domain (residues 209-582) and UBL1-5 domain (residues 583-1192) of USP7 did not (Fig. 2D). Deletion of the TRAF domain significantly impaired the interaction between PAF15 and USP7, although it did not affect the binding to DNMT1. This is further supported by the observation that the introduction of the W167A mutation, which disrupts the TRAF binding pocket (Sheng et al., 2006), resulted in a loss of interaction with PAF15. Deletion of the UBL domain or mutations into the UBL2 pocket, D758A/E759A/D764A (Cheng, Yang, et al., 2015), also decreased binding to PAF15. Since the UBL1-5 domain alone is not sufficient for PAF15 binding, this interaction might only occur in the context of the full-length protein. Taken together, these results indicate that both the TRAF and UBL2 domains of USP7 contribute to the interaction with PAF15, as has been recently reported for other USP7 substrates (Ashton et al., 2021; Georges et al., 2019).

### USP7 is involved in PAF15 dissociation from chromatin during S phase progression

We next tested whether USP7 regulates PAF15 on chromatin. To this end, we examined the chromatin binding of a recombinant PAF15 mutant lacking USP7 binding activity in PAF15-depleted extracts. As seen for endogenous PAF15, wild-type rPAF15 dissociated from chromatin at 120 min when added to PAF15-depleted egg extracts (Fig. 3A, S2A). In contrast, the rPAF15 with mutated USP7 interacting sequences (SAKA) showed prolonged chromatin association even after 120 min, although chromatin binding of USP7 was not affected. To directly test the importance of USP7 for regulation of PAF15 chromatin binding, we depleted USP7 from egg extracts. Compared to the control, the USP7-depleted extracts showed impaired dissociation of PAF15 chromatin (Fig. 3B, S2B). Affinity-purified recombinant USP7 efficiently restored PAF15 chromatin dissociation, but the USP7 C225S, C223 in human, catalytic inactive mutant failed to do so (Hu et al., 2002; Li et al., 2002) (Fig. 3B, S2B). The USP7 specific inhibitor FT671 also suppressed the PAF15Ub2 chromatin dissociation (Fig. 3C, S2C). These results suggested that USP7 regulates PAF15 chromatin dissociation through its deubiquitylation activity. To investigate whether USP7 directly deubiquitylates PAF15, we performed an *in vitro* deubiquitylation assay by using purified recombinant ubiquitylated hPAF15 and hUSP7. We ubiquitylated hPAF15 by incubating with E1 (mouseUBA1), E2 (UBE2D3), E3 (UHRF1) enzymes *in vitro*. After purification, we incubated ubiquitylated hPAF15 with recombinant hUSP7 and analyzed the reaction products. USP7 WT efficiently deubiquitylated PAF15 while the catalytic inactive USP7 mutant (C223A) did not (Fig. 3D). USP47 that is closely related DUB to USP7 did not show deubiquitylation activity toward the ubiquitylated PAF15 (Fig. S2D). These results suggested that USP7 directly deubiquitylates PAF15 to promote PAF15 chromatin dissociation.

**Figure 3.**
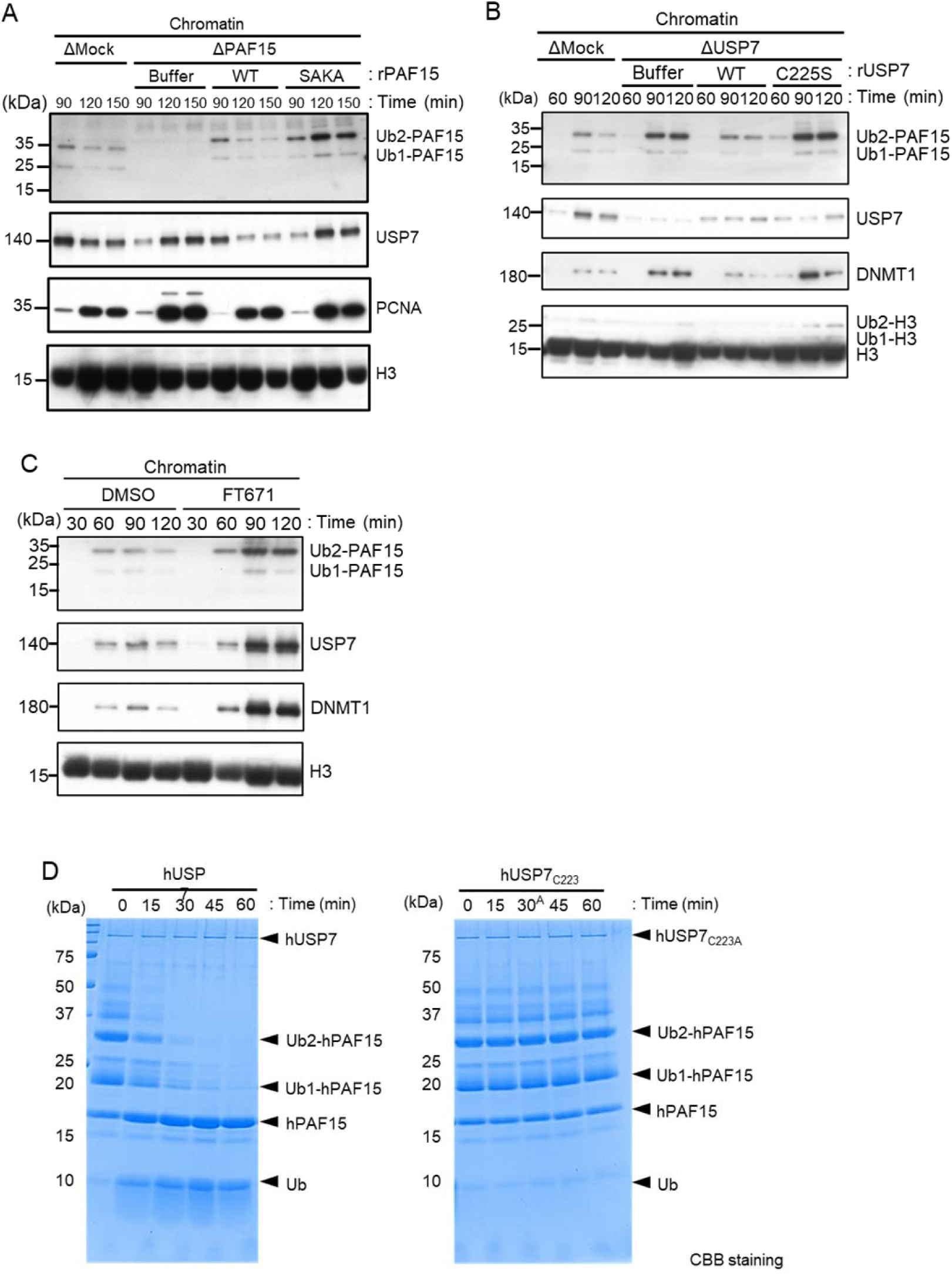
USP7 promotes PAF15 dissociation from chromatin. **(A)** Sperm chromatin was added to Mock- or PAF15-depleted interphase extracts supplemented with either buffer (+Buffer), wild-type rPAF15-3xFLAG (+WT) or rPAF15 SAKA-3xFLAG (+SAKA). Chromatin fractions were isolated, and the samples were analyzed by immunoblotting using the antibodies indicated. **(B)** Sperm chromatin was added to Mock- or USP7-depleted interphase extracts supplemented with either buffer (+Buffer), wild-type rUSP7-3xFLAG (+WT) or catalytic mutant rUSP7 C225S-3xFLAG (+C225S). Chromatin fractions were isolated, and the samples were analyzed by immunoblotting using the antibodies indicated. (C) Sperm chromatin was added to interphase extracts supplemented with either DMSO (+DMSO), or FT671 (+FT671). Chromatin fractions were isolated, and the samples were analyzed by immunoblotting using the antibodies indicated. (D) Ubiquitylated hPAF15 was incubated with recombinant hUSP7 WT (left) or C223A catalytic mutant (right) at indicated times. The reaction products were analyzed by SDS-PAGE with CBB staining. Source Data are provided as Figure 3-source data.

To further elucidate the mechanism underlying termination of PAF15 signaling by USP7, we examined whether USP7 interacts with PAF15 on chromatin. As shown in previous reports (Nishiyama et al., 2020), chromatin-bound PAF15 existed mainly as ubiquitylated forms (PAF15Ub2 or PAF15Ub1), and PAF15Ub2 specifically interacted with DNMT1 (Fig. 4A). USP7 co-immunoprecipitated with PAF15 as well as DNMT1. Importantly, PAF15Ub2 was readily detected in the USP7 immunoprecipitates, whereas PAF15Ub1 and PAF15Ub0 were not. Given that DNMT1 forms a complex with USP7 and predominantly binds to PAF15Ub2, USP7 binding to PAF15 might be mediated by DNMT1 at DNA methylation sites.

**Figure 4.**
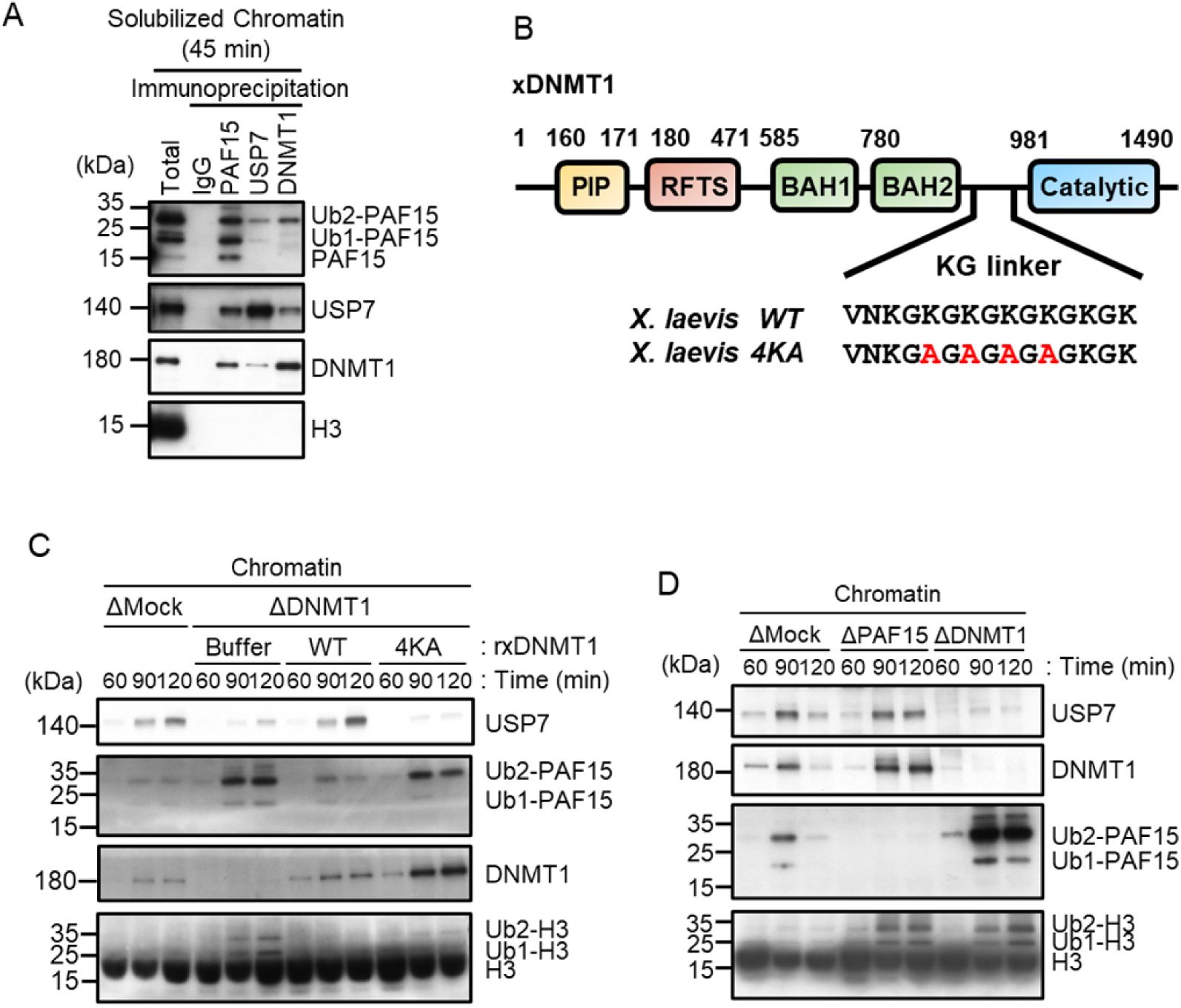
USP7 is recruited to chromatin through the interaction with DNMT1 for PAF15 deubiquitylation. **(A)** Sperm chromatin was added to interphase extracts. Replicating chromatin was digested by micrococcal nuclease (MNase). Immunoprecipitation was performed by PAF15, USP7 and DNMT1 antibodies from the solubilized chromatin fraction, and the samples were analyzed by immunoblotting using the antibodies indicated. **(B)** Illustration of DNMT1 domain structure. The KG linker located between the BAH domain and catalytic domain contributes to interaction with USP7. rDNMT1 4KA mutant, using in (C), was introduced mutation at four lysines to alanine in the KG linker. **(C)** Sperm chromatin was added to Mock- or DNMT1-depleted interphase extracts supplemented with either buffer (+Buffer), wild-type rDNMT1-3xFLAG (+WT), or rDNMT1 4KA-3xFLAG (+4KA). Chromatin fractions were isolated, and the samples were analyzed by immunoblotting using the antibodies indicated. **(D)** Sperm chromatin was added to Mock-, PAF15-, and DNMT1-depleted extracts. Chromatin fractions were isolated, and the samples were analyzed by immunoblotting using the antibodies indicated. Source Data are provided as Figure 4-source data.

In order to inhibit the interaction between DNMT1 and USP7, we introduced mutations into the KG linker of DNMT1, which is responsible for USP7 binding (Cheng, yang, et al., 2015; Yamaguchi et al., 2017). We performed immunodepletion of endogenous DNMT1 from egg extracts and added-back wild-type DNMT1 or the KG linker mutant (DNMT1 4KA). As previously reported, immunodepletion of DNMT1 inhibited the chromatin binding of USP7 and induced marked accumulation of ubiquitylated PAF15 and histone H3 (Nishiyama et al., 2013; Nishiyama et al., 2020). Wild-type DNMT1 efficiently restored USP7 chromatin recruitment and PAF15 chromatin dissociation, but DNMT1 4KA failed to do so (Fig. 4B, C, S3A). These results suggest that DNMT1 recruits USP7 to chromatin and mediates the formation of USP7-PAF15Ub2 complex. Consistent with this idea, immunodepletion of PAF15 had no significant effect on the level of chromatin-bound USP7 (Fig. 4D, S3B).

### Unloading of PAF15 couples with the completion of DNA methylation maintenance

We next tested how PAF15 chromatin dissociation is coordinated with the progression of maintenance DNA methylation. Completion of DNA maintenance methylation is accompanied by conversion of hemimethylated DNA to fully methylated DNA by DNMT1 and subsequent inactivation of UHRF1-dependent ubiquitin signaling. To inhibit DNMT1 activity, we replaced the endogenous DNMT1 with a recombinant DNMT1 C1101S mutant that lacks DNMT1 catalytic activity (Takebayashi et al., 2007; Takeshita et al., 2011; Wyszynski et al., 1993). The results showed that the inactivation of DNMT1 led to accumulation of UHRF1 on chromatin, presumably due to the failure in the conversion of hemi-methylated DNA to fully methylated state (Fig. 5A, S4A). PAF15Ub2 showed a significant accumulation on chromatin along with H3Ub2 under this condition, suggesting that the completion of maintenance DNA methylation is required for the USP7-dependent dissociation of PAF15 from chromatin. Note that the recruitment of USP7 to chromatin was rather enhanced when DNMT1 was inactivated (Fig. 5A). Immunoprecipitation of PAF15 or USP7 from chromatin lysates showed that inhibition of the catalytic activity of DNMT1 did not affect the binding of USP7 to PAF15Ub2 (Fig. 5B). These results suggest that the USP7-mediated deubiquitylation couples the completion of DNA methylation by DNMT1.

**Figure 5.**
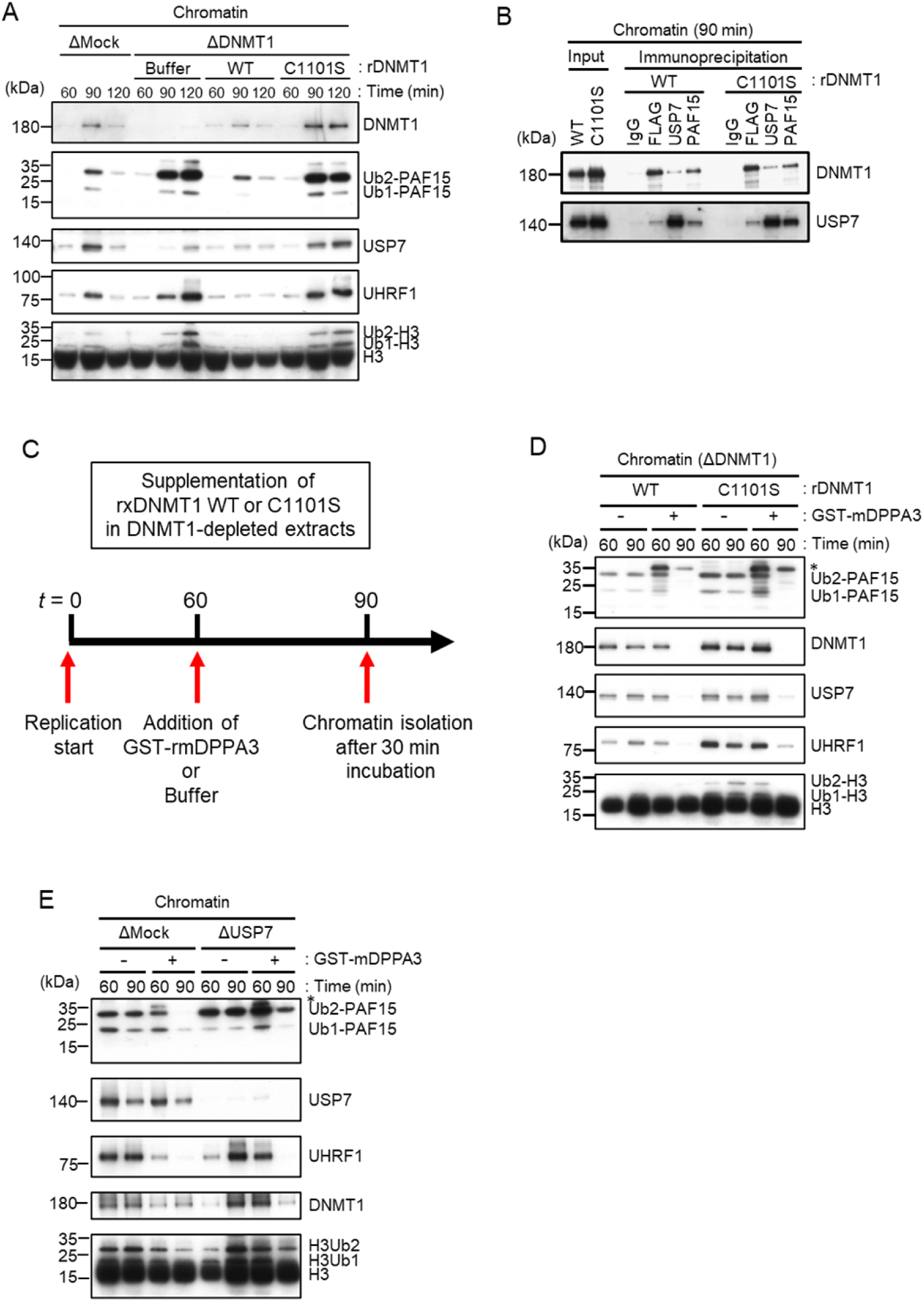
Unloading of PAF15 requires DNMT1-dependent DNA methylation. **(A)** Sperm chromatin was added to Mock- or DNMT1-depleted interphase extracts supplemented with either buffer (+Buffer), wild-type rDNMT1-3xFLAG (+WT), or catalytic mutant rDNMT1 C1101S-3xFLAG (+C1101S). Chromatin fractions were isolated, and the samples were analyzed by immunoblotting using the antibodies indicated. **(B)** Sperm chromatin was added to DNMT1-depleted interphase extracts supplemented wild-type rDNMT1-3xFLAG (+WT) or catalytic mutant rDNMT1 C1101S-3xFLAG (+C1101S). Replicating chromatin was digested by MNase. Immunoprecipitation was performed by PAF15, USP7 antibodies-bound beads, and FLAG affinity beads in the solubilized chromatin fraction solution, and the samples were analyzed by immunoblotting using the antibodies indicated. **(C)** A schema of an experiment described in Figure 5C. **(D)** Sperm chromatin was added to DNMT1-depleted interphase extracts supplemented wild-type rDNMT1-3xFLAG (+WT) or catalytic mutant rDNMT1 C1101S-3xFLAG (+C1101S). After 60 min, the extracts were supplemented with either buffer (-) or GST-mDPPA3 61-150 (+). Chromatin fractions were isolated, and the samples were analyzed by immunoblotting using the antibodies indicated. The asterisk indicates a non-specific band. **(E)** Sperm chromatin was added to USP7-depleted interphase extracts. After 90 min, the extracts were supplemented with either buffer (-) or GST-mDPPA3 61-150 (+). Chromatin fractions were isolated, and the samples were analyzed by immunoblotting using the antibodies indicated. The asterisk indicates a non-specific band. Source Data are provided as Figure 5-source data.

Failure of DNA methylation replication has been shown to be accompanied by accumulation of UHRF1 on chromatin and enhanced UHRF1-dependent ubiquitin signaling. We hypothesized that when maintenance DNA methylation is inhibited, the enhanced E3 ligase activity of UHRF1 caused by its chromatin accumulation may overcome the deubiquitylation activity of USP7, which apparently suppresses the deubiquitylation of PAF15. Recent studies have reported that the maternal gene Stella/DPPA3, which protects against oocyte-specific DNA methylation in mice, binds directly to the UHRF1-PHD domain and inhibits UHRF1 nuclear localization and chromatin binding (Du et al., 2019; Li et al., 2018; Mulholland et al., 2020). We have previously shown that the addition of recombinant mouse DPPA3 to egg extracts inhibits the chromatin-binding activity of UHRF1 and induces dissociation of UHRF1. To determine whether UHRF1 competes with the deubiquitylation by USP7, we forced chromatin dissociation of UHRF1 by adding the purified recombinant GST-mDPPA3 to DNMT1 depleted extracts supplemented with the DNMT1 C1101S mutant (Fig. 5C). The addition of recombinant mDPPA3 efficiently induced chromatin dissociation of UHRF1, leading to a significant decrease in the levels of chromatin-bound PAF15 and DNMT1 (Fig. 5D, S4B). Importantly, USP7 depletion caused significant delay of PAF15 chromatin dissociation induced by mDPPA3 addition. These results suggest that UHRF1 maintains PAF15 chromatin association by counteracting USP7-dependent PAF15 deubiquitylation until the completion of maintenance DNA methylation.

### ATAD5 promotes PAF15 unloading from chromatin

It has been shown that PCNA is unloaded from chromatin by the ATAD5-RLC (RFC-like complex) in a coordinated manner with the maturation of Okazaki fragment in the late S phase (Johnson et al., 2016; Kang et al., 2019; Kubota et al., 2015; Kubota et al., 2013; Lee et al., 2013; Ulrich, 2013). Based on the requirement of PCNA for PAF15 chromatin loading described previously, we investigated the role of ATAD5 in the chromatin dissociation of PAF15. Consistent with previous reports in mammalian cultured cells and budding yeast, immunodepletion of ATAD5 from interphase egg extracts resulted in the chromatin accumulation of PCNA (Fig. 6A) (Kubota et al., 2013; Lee et al., 2013). Interestingly, chromatin binding of PAF15Ub0 was readily detected on ATAD5-depleted chromatin (Fig. 6A, S5A). On the other hand, no significant change was observed in the amount of PAF15Ub2 on chromatin in ATAD5-depleted extracts. In USP7/ATAD5 co-depleted extracts, PAF15 showed accumulation on chromatin regardless of its ubiquitylation state. The accumulation of the PAF15Ub0 and Ub1 were rescued by recombinant hATAD5-RFCs addition to ATAD5-depleted extracts (Fig. 6B, S5B). These results suggested that ATAD5 regulates PAF15Ub0 and Ub1 chromatin dissociation.

**Figure 6.**
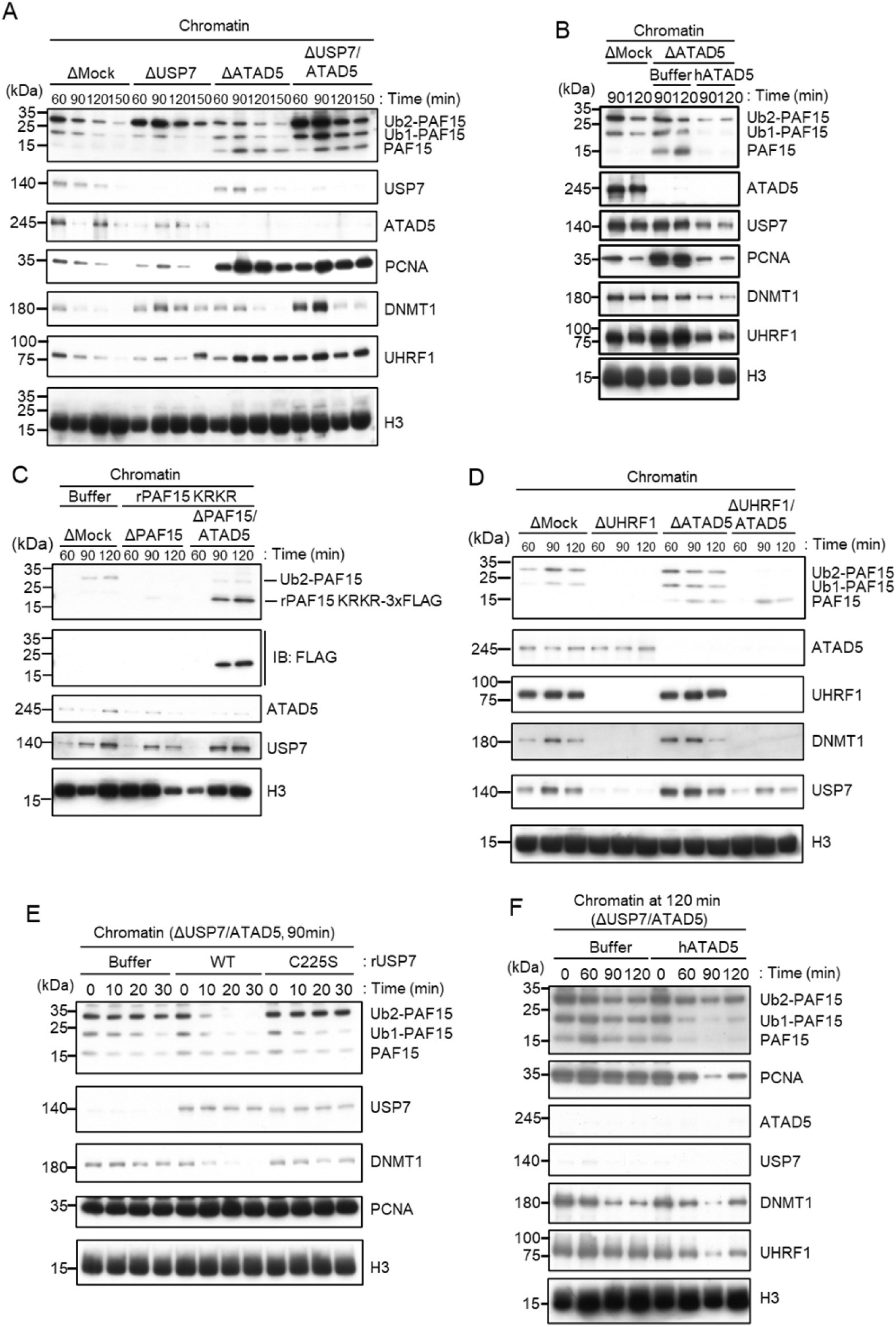
ATAD5 unloads PAF15Ub0 from chromatin. **(A)** Sperm chromatin was added to Mock-, USP7-, ATAD5-, and USP7/ATAD5-depleted interphase extracts, and isolated at indicated time points. Chromatin bound proteins were analyzed by immunoblotting. (B) Sperm chromatin was added to Mock-, ATAD5-depleted interphase extracts supplemented with either buffer (+Buffer) or recombinant hATAD5-RFCs (+ATAD5). Chromatin fractions were isolated, and the samples were analyzed by immunoblotting using the antibodies indicated. **(C)** Recombinant PAF15 K18R/K27R-3xFLAG was supplemented to PAF15-, and PAF15/ATAD5-depleted extracts, and chromatin fractions were isolated. Chromatin bound proteins were confirmed by immunoblotting. **(D)** Sperm chromatin was added to Mock-, UHRF1-, ATAD5-, and UHRF1/ATAD5-depleted extracts, and isolated at indicated time point. Chromatin bound proteins were analyzed by immunoblotting. **(E)** Sperm chromatin was added to USP7/ATAD5-depleted extracts, and isolated at 90 min. The chromatin was supplemented with either buffer (+Buffer), USP7 WT-3xFLAG (+WT), or USP7 C225S-3xFLAG (+C225S) and re-isolated at indicated time points. Chromatin bound proteins were analyzed by immunoblotting. **(F)** Sperm chromatin was added to Mock-, USP7/ATAD5-depleted extracts. After replication at 90 min, the extracts were supplemented with either buffer (+Buffer) or recombinant hATAD5-RFCs (+ATAD5). Chromatin fractions were isolated and the samples were analyzed by immunoblotting using the antibodies indicated. Source Data are provided as Figure 6-source data.

We next examined whether ATAD5 regulates chromatin unloading of non-ubiquitylated PAF15. First, we added a PAF15 mutant lacking the ubiquitylation sites (PAF15 KRKR) to the PAF15/ATAD5 double-depleted extract and examined its chromatin binding. As expected, PAF15 KRKR did not show chromatin binding in the presence of ATAD5, but its chromatin binding became detectable in ATAD5-depleted extracts (Fig. 6C, S5C). We also inhibited PAF15 ubiquitylation by UHRF1 depletion. As previously reported, UHRF1 depletion completely inhibited PAF15 ubiquitylation and chromatin loading, resulting in inhibition of DNMT1 recruitment (Nishiyama et al., 2020). However, obvious chromatin binding of non-ubiquitylated PAF15 was observed in UHRF1/ATAD5 double-depleted extracts (Fig. 6D, S5D). These results suggest that non-ubiquitylated or deubiquitylated PAF15 is unloaded in an ATAD5-dependent manner.

### USP7 and ATAD5 promote dissociation of chromatin-bound PAF15

Next, we investigated whether USP7 and ATAD5 accelerates PAF15 chromatin dissociation. To induce PAF15 chromatin accumulation, sperm chromatin was incubated in USP7/ATAD5 co-depleted extracts. Chromatin was isolated after 90 min replication and further incubated with recombinant USP7 WT or C225S catalytic inactive mutant. PAF15Ub2, but not PAF15Ub1 or PAF15Ub0, was efficiently dissociated from chromatin only when recombinant wild-type USP7 was added (Fig. 6E, S5E). Conversely, when recombinant hATAD5-RLCs was added to USP7/ATAD5 co-depleted extracts after 120 min replication, PAF15Ub1 and Ub0 dissociated from chromatin (Fig. 6F, S5F). These results suggested that chromatin dissociation of PAF15Ub2 is regulated by USP7, whereas PAF15Ub1 and PAF15Ub0 are regulated by ATAD5-dependent unloading.

### Co-depletion of USP7 and ATAD5 leads to an increase in global DNA methylation

Inhibition of chromatin unloading of PAF15 might affect maintenance DNA methylation. To investigate changes in the level of global DNA methylation and efficiency of DNA replication by USP7- and/or ATAD5-depletion, we measured the incorporation of radiolabeled S-adenosyl-methionine (^3^H-SAM) and [α-^32^P] dCTP into DNA, respectively. USP7/ATAD5 double-depletion caused increased DNA methylation compared to mock-depleted extracts without significant effect on gross DNA replication (Fig. 7, S6A, B), but either USP7- or ATAD5-depletion alone did not do so. These data suggest that the termination of PAF15 ubiquitin signaling suppresses excessive DNA methylation.

**Figure 7.**
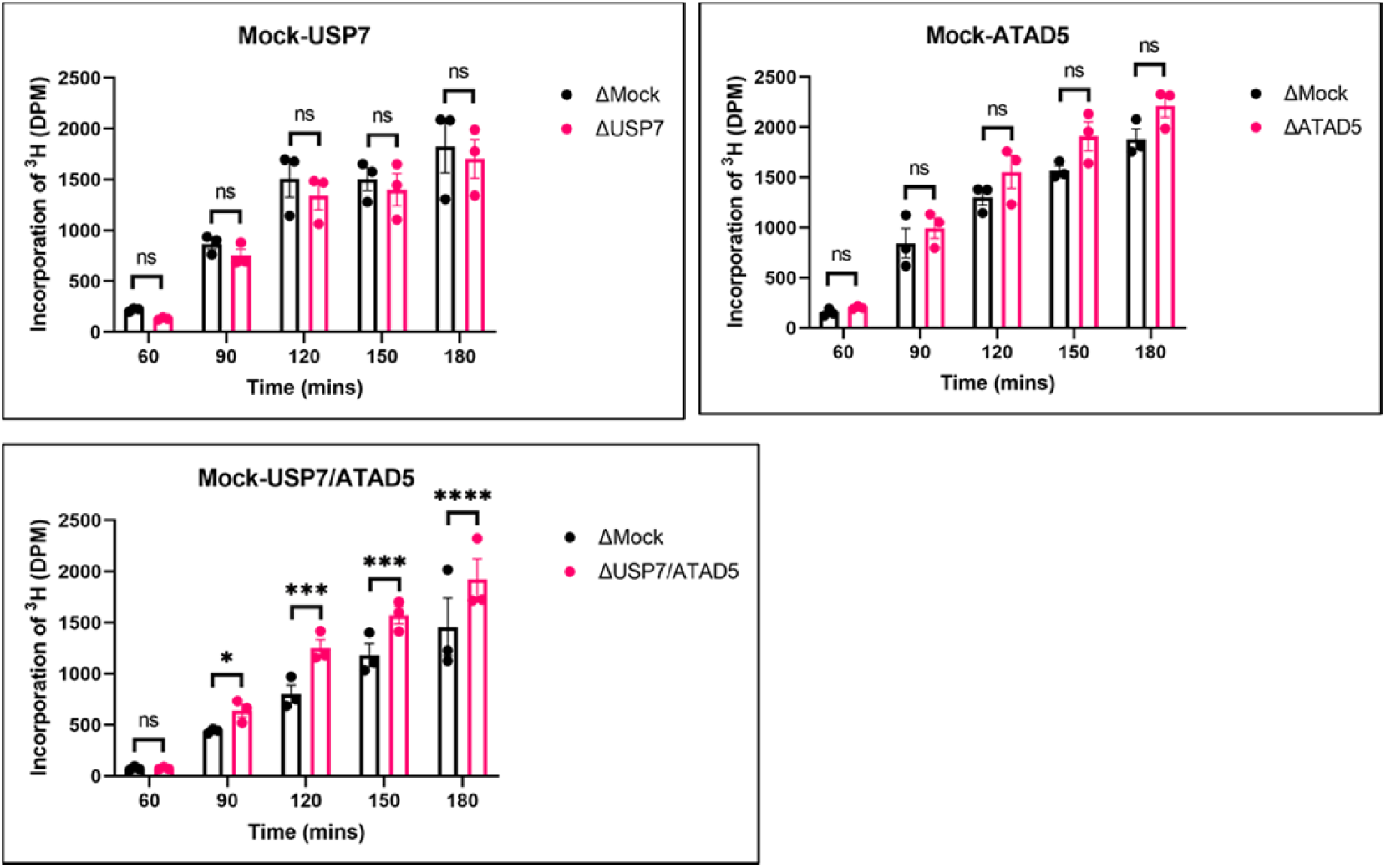
PAF15 dissociation negatively regulates aberrant increase of DNA methylation. Sperm chromatin and radiolabeled S-[methyl-^3^H]-adenosyl-L-methionine were added to either Mock- and USP7-, ATAD5-, or USP7/ATAD5 co-depleted extracts. Purified DNA samples were analyzed to determine the efficiency of DNA methylation. Data are presented as mean ± SEM. Multiple comparisons were performed by Two-way Repeated Measure analysis of variance (RM ANOVA) followed by Sidak’s multiple comparison test. ns; not significant, ∗p < 0.05, ∗∗∗p < 0.001, ∗∗∗∗p < 0.0001. Source Data are provided as Figure 7-source data.

**Figure 8.**
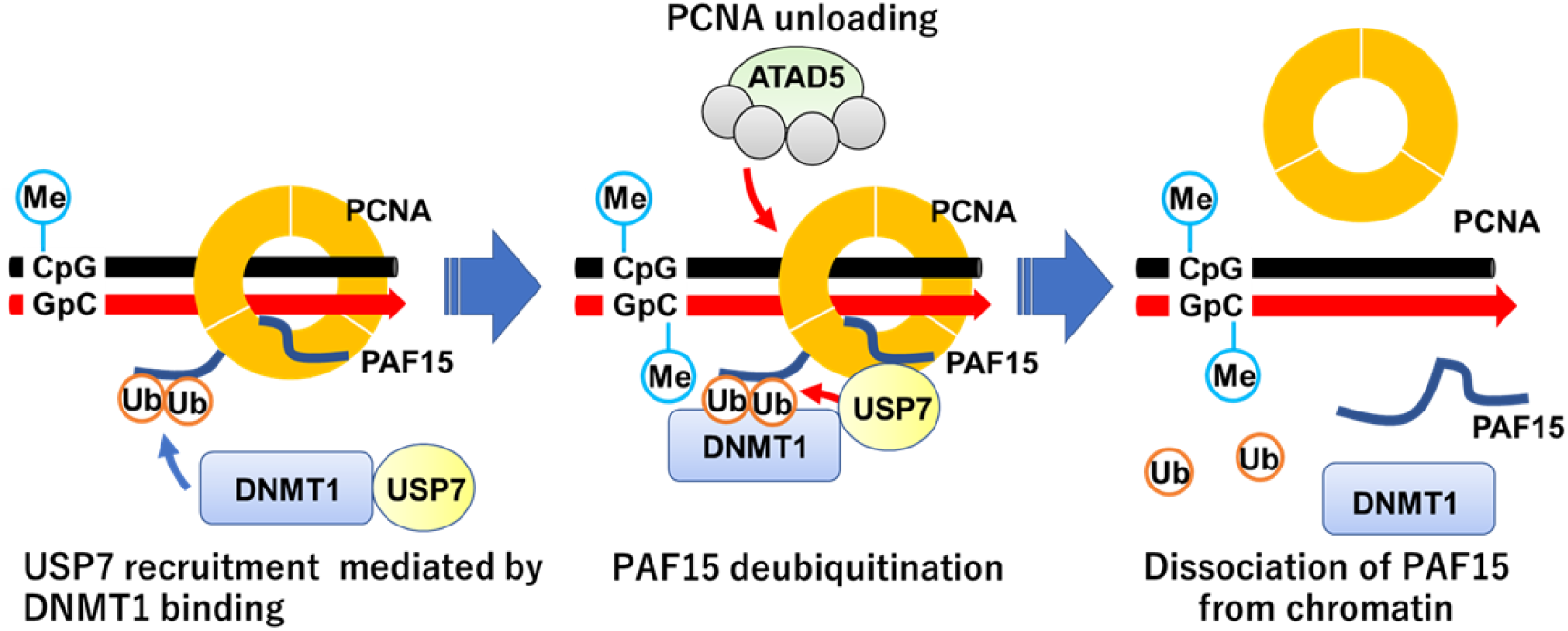
A schematic model of PAF15 deubiquitylation and chromatin unloading During the S phase, USP7 is recruited to the methylation site along with DNMT1. After the conversion of hemi-methylated DNA to fully methylated DNA by DNMT1, USP7 promotes deubiquitylation of PAF15Ub2. Subsequently, the deubiquitylated PAF15 (PAF15Ub0) is removed from chromatin together with PCNA by the ATAD5-RLC complex.

## Discussion

In this study, we investigated how PAF15 chromatin unloading is regulated during the completion of maintenance DNA methylation. Using *Xenopus* egg extracts, we demonstrate that PAF15 unloading is regulated by two distinct mechanisms. First, USP7 deubiquitylates PAF15Ub2 to promote PAF15Ub2 dissociation from chromatin. Second, the ATAD5-RLCs complex promotes chromatin unloading of non-ubiquitylated PAF15 and PAF15Ub1 together with PCNA. Importantly, our data show that co-depletion of USP7 and ATAD5 leads to chromatin accumulation of DNMT1 together with PAF15Ub2 and increased global DNA methylation. Consistent with this data, previous report also showed that the loss of USP7 in HeLa cells leads to the increase of DNA methylation in a substantial fraction of de novo DNA methylation sites upon long-term culture (Li et al., 2020). We speculate that accumulation of both PAF15Ub2 and PCNA on USP7/ATAD5-depleted chromatin causes premature DNMT1 localization and hyperactivation at de novo DNA methylation sites.

Our data showed that USP7 specifically targets PAF15Ub2 to facilitate its chromatin dissociation; PAF15Ub2 has been reported to interact with the RFTS domain of DNMT1 (Nishiyama et al., 2020), thereby recruiting the DNMT1-USP7 complex to the methylation sites. Since the interaction between USP7 and PAF15 in the cell is probably very weak, the formation of a stable recognition of PAF15Ub2 by USP7. Notably, neither removal nor inhibition of USP7 enhanced histone H3 ubiquitylation, suggesting that PAF15 ubiquitin signaling is the primary pathway to maintain DNA methylation during S phase as previously reported (Nishiyama et al., 2020). Although our results indicate that USP7 functions as the major DUB for PAF15 deubiquitylation, it remains possible that other DUBs also influence this process. Indeed, the gradual decrease in chromatin binding of PAF15Ub2 in the absence of USP7 suggests that other proteins are also involved.

Our results also demonstrate that the ATAD5-RLC complex is required for chromatin dissociation of non-ubiquitylated PAF15 and PAF15Ub1. Notably, many of DNA replication proteins interacting with PCNA compete for binding surfaces between ATAD5 and PCNA during DNA replication (Kang et al., 2019). We speculate that PAF15 may not interfere with the interaction between ATAD5 and PCNA due to its small size and flexible structure. Our results suggested that PAF15Ub2 is not targeted by ATAD5-dependent unloading. This is consistent with our previous report that dual mono-ubiquitylation of PAF15 plays a pivotal role in its chromatin binding (Nishiyama et al., 2020). However, it is not clear how dual mono-ubiquitylation of PAF15 contributes to stable PAF15 chromatin binding. Interestingly, several studies reported the DNA binding activity of ubiquitin. K63-linked ubiquitin chain binds to DNA directly through its DNA interacting patch, consists with threonine, lysine, and glutamic acid (Liu et al., 2018). Another paper also has shown that mono-ubiquitylation of transcription factors, such as p53 or IRF1 enhanced their nuclear localization and chromatin binding (Landre et al., 2017). Future biochemical analyses will be required to test whether dual monoubiquitylation enhances DNA binding activity of PAF15. Alternatively, DNMT1, which forms a complex with PAF15Ub2, may prevent ATAD5-dependent unloading by interacting with PCNA via the PIP-box. Intriguingly, the chromatin binding of non-ubiquitylated PAF15 in ATAD5-depleted extracts did not require UHRF1. These data suggest that PCNA-mediated loading of PAF15 could occur at sites outside DNA methylation sites. In such regions, ATAD5 may prevent the formation of the maintenance DNA methylation machinery by excluding the PAF15-PCNA complex from chromatin.

In summary, data presented here suggest that the coupled ubiquitylation and deubiquitylation may be necessary for proper maintenance of DNA methylation. Interestingly, inactivation of DNMT1 catalytic activity almost completely suppressed chromatin dissociation of PAF15. How is the PAF15 inactivation coupled to the completion of methylation for DNA maintenance? Even when DNA methylation was inhibited, chromatin recruitment of USP7 and the formation of the USP7-PAF15Ub2 complex were observed. Thus, the inhibition of deubiquitylation in this condition is not caused by suppressing USP7 chromatin recruitment or USP7 interaction with PAF15Ub2. Our results suggest that ubiquitylation by UHRF1 is predominant over the deubiquitylation and unloading of PAF15, maintaining PAF15Ub2 until the completion of maintenance methylation by DNMT1. UHRF1 is thought to dissociate from DNA upon binding to hemi-methylated DNA by DNMT1 (Arita et al., 2008). Dissociation of UHRF1 from chromatin upon the conversion of hemi-methylated DNA to fully methylated DNA would trigger the removal of ubiquitin moieties from PAF15 by USP7. It is also possible that the binding of DNMT1 to hemi-methylated DNA induces conformational changes in USP7 or PAF15Ub2 to facilitate the deubiquitylation of PAF15Ub2 by USP7. Detailed analysis of the DNMT1-USP7-PAF15Ub2 complex will be important in future studies.

## Methods

### Primers

All oligonucleotide sequences are listed in the Supplementary Table 3.

### *Xenopus* egg extracts

*Xenopus laevis* was purchased from Kato-S Kagaku and handled according to the animal care regulations at the University of Tokyo. The preparation of interphase egg extracts, chromatin isolations, UbVS reactions, DNA replication assays, DNA methylation assays, and immunodepletions were performed as described previously (Kumamoto et al., 2021; Nishiyama et al., 2020). Unfertilized *Xenopus laevis* eggs were dejellied in 2.5 % thioglycolic acid-NaOH (pH 8.2) and washed in 1xMMR buffer [100 mM NaCl, 2 mM KCl, 1 mM MgCl_2_, 2 mM CaCl_2_, 0.1 mM Ethylenediaminetetraacetic acid (EDTA), 5 mM HEPES-NaOH (pH 7.5)]. After activation in 1xMMR supplemented with 0.3 µg/ml calcium ionophore, eggs were washed with EB buffer [50 mM KCl, 2.5 mM MgCl_2_, 10 mM HEPES-KOH (pH 7.5), 50 mM sucrose]. Eggs were packed into tubes by centrifugation (BECKMAN, Avanti J-E, JS-13.1 swinging rotor) for 1 min at 190 xg and crushed by centrifugation for 20 min at 18,973 xg. Egg extracts were supplemented with 50 µg/mL cycloheximide, 20 µg/mL cytochalasin B, 1 mM dithiothreitol (DTT), 2 µg/mL aprotinin, 5µg/mL leupeptin and clarified by ultracentrifugation (Hitachi, CP100NX, P55ST2 swinging rotor) for 20 min at 48,400 xg. The cytoplasmic extracts were aliquoted, frozen in liquid nitrogen, and stored at -80 °C. All extracts were supplemented with an energy regeneration system (2 mM ATP, 20 mM phosphocreatine, and 5 µg/ml creatine phosphokinase). 3,000-4,000 nuclei/µl of sperm nuclei were added and incubated at 22 °C. Aliquots (15-20 µl) were diluted with 150 µl chromatin purification buffer [CPB; 50 mM KCl, 5 mM MgCl_2_, 20 mM HEPES-KOH (pH 7.6)] containing 0.1% Nonidet P-40 (NP-40), 2% sucrose, 2 mM N-ethylmaleimide (NEM) and 0.1 mM PR-619. After incubation on ice for 5 min, diluted extracts were layered over 1.5 ml of CPB containing 30% sucrose and centrifuged at 15,000 xg for 10 min at 4 °C. Chromatin pellets were resuspended in 1xLaemmli sample buffer, boiled for 5 min at 100 °C, and analyzed by immunoblotting.

### Antibodies and immunoprecipitations/immunodepletions

*Xenopus* ATAD5 (xATAD5) antibodies were raised in rabbits by immunization with a His10-tagged recombinant xATAD5 fragment encoding 1-289 amino acids and used for immunodepletion and immunoblotting. Rabbit polyclonal antibodies raised against PAF15, DNMT1, and UHRF1 have been previously described. Rabbit polyclonal USP7 antibody (A300-033A) was purchased from Bethyl Laboratories. Mouse monoclonal antibody against PCNA (PC-10) was purchased from Santa Cruz Biotechnology. Rabbit polyclonal histone H3 antibody (ab1791) was purchased from Abcam. For immunoprecipitation, 10 µl of Protein A agarose (GE Healthcare) was coupled with 2 µg of purified antibodies or 5 µl of antiserum. The agarose beads were washed twice with CPB buffer containing 2% sucrose. The antibody beads were incubated with egg extracts for 2 h at 4 °C. The beads were washed three times with CPB buffer containing 2% sucrose and 0.1% Triton X-100 and resuspended in 10 µl of 2xLaemmli sample buffer and 20 µl of 1xLaemmli sample buffer. For immunodepletion, 250 µl of antiserum were coupled to 60 µl of recombinant Protein A Sepharose (rPAS, GE Healthcare). Antibodies bound beads were washed three times in CPB and supplemented with 6 µl fresh rPAS. Beads were split into three portions, and 100 µl of extracts were depleted in three rounds at 4 °C, each for 1 h.

### GST pull-down assay in *Xenopus* egg extracts

Recombinant GST or GST-PAF15 proteins were expressed and purified from *Escherichia. Coli* (BL21-CodonPlus), and immobilized on Glutathione Sepharose 4B resin (GE Healthcare) for 2 h at 4 °C. The beads were incubated with interphase egg extracts for 2 h at 4 °C. The beads were washed four times with CPB containing 2% sucrose and 0.1% Triton X-100. The washed beads were resuspended in 20 µl of 2xLaemmli sample buffer and 20 µl of 1xLaemmli sample buffer, boiled for 5 min at 100 °C, and analyzed by immunoblotting.

### Immunoprecipitation of FLAG-USP7

Recombinant 3xFLAG-tagged USP7 proteins were expressed in Sf9 insect cells. These insect cells were collected and suspended in lysis buffer [20 mM Tris-HCl (pH 8.0), 100 mM KCl, 5 mM MgCl_2_, 10% glycerol, 1% NP-40, 1 mM DTT, 5 µg/ml leupeptin, 2 µg/ml aprotinin, 20 µg/ml trypsin inhibitor, 100 µg/ml phenylmethylsulfonyl fluoride (PMSF)], followed by incubation on ice for 10 min. Soluble fractions were isolated after centrifugation of the lysate at 15,000 xg for 15 min at 4 °C. 2 ml of the soluble lysate was incubated with 30 µl of anti-FLAG M2 affinity resins (Sigma-Aldrich) for 2 h at 4 °C. The protein-bound beads were washed five times with wash buffer [20 mM Tris-HCl (pH 8.0), 100 mM KCl, 5 mM MgCl_2_, 10% glycerol, 0.1% NP-40, 1 mM DTT, 5 µg/ml leupeptin, 2 µg/ml aprotinin, 20 µg/ml trypsin inhibitor, 100 µg/ml PMSF] and stored in PBS at 4 °C. 10 µl of protein-bound FLAG beads were coupled with 100 µl of *Xenopus* egg extracts diluted five-fold by CPB containing 2% sucrose and incubated for 2 h at 4 °C. The beads were washed three times by CPB containing 2% sucrose and 0.1% Triton X-100, followed by resuspension by 10 µl of 2xLaemmli sample buffer and 20 µl of 1xLaemmli sample buffer.

### Mass spectrometry

The eluted proteins were trypsin-digested, desalted using ZipTip C18 (Millipore) and centrifuged in a vacuum concentrator. Shotgun proteomic analyses of the digested peptides were performed by LTQ-Orbitrap Velos mass spectrometer (Thermo Fisher Scientific) coupled with Dina-2A nanoflow liquid chromatography system (KYA Technologies). The samples were injected into a 75-µm reversed-phase C18 column at a flow rate of 10 µl/min and eluted with a linear gradient of solvent A (2% acetonitrile and 0.1% formic acid in H_2_O) to solvent B (40% acetonitrile and 0.1% formic acid in H_2_O) at 300 nl/min. Peptides were sequentially sprayed from a nanoelectrospray ion source (KYA Technologies) and analyzed by collision-induced dissociation (CID). The analyses were operated in data-dependent mode, switching automatically between MS and MS/MS acquisition. For CID analyses, full-scan MS spectra (from m/z 380 to 2,000) were acquired in the orbitrap with a resolution of 100,000 at m/z 400 after ion count accumulation to the target value of 1,000,000. The 20 most intense ions at a threshold above 2,000 were fragmented in the linear ion trap with a normalized collision energy of 35% for an activation time of 10 ms. The orbitrap analyzer was operated with the “lock mass” option to perform shotgun detection with high accuracy. Protein identification was conducted by searching MS and MS/MS data against NCBI (National Center for Biotechnology Information) *Xenopus laevis* protein database using Mascot (Matrix Science). Methionine oxidation, protein N-terminal acetylation and pyro-glutamination for N-terminal glutamine were set as variable modifications.

A maximum of two missed cleavages was allowed in our database search, while the mass tolerance was set to three parts per million (ppm) for peptide masses and 0.8 Da for MS/MS peaks. In the process of peptide identification, we applied a filter to satisfy a false discovery rate lower than 1%. The mass spectrometry proteomics data have been deposited to the ProteomeXchange Consortium via the jPOST repository with the dataset identifier PXD034088.

### *In vitro* deubiquitylation assay

Ubiquitylated PAF15, a substrate for the deubiquitylation assay, was prepared by *in vitro* ubiquitylation using recombinant mouse UBA1 (E1), human UBE2D3 (E2), human UHRF1 (E3), ubiquitin and PAF15. N-terminal six histidine tagged E1 was expressed in Sf9 cells using the baculo virus system according to the manufacture’s instruction. The protein was purified by TALON affinity (Clontech), HiTrap-Q anion-exchange (Cytiva) and Hiload 26/600 S200 size-exclusion (Cytiva) chromatographies. E2 was expressed in *Escherichia coli* BL21 as a GST-fusion protein and purified by GS4B affinity (Cytiva) and d Hiload 26/600 S75 size-exclusion chromatographies (Cytiva). UHRF1 was expressed in *Escherichia coli* Rossetta2 and purified using GS4B affinity, HiTrap Heparin and Hiload 26/600 S200 size-exclusion chromatographies. Ubiquitin was expressed in BL21 and purified using HiTrap SP anion-exchange (Cytiva) and Hiload 26/600 S75 size-exclusion chromatographies. PAF15 including C-terminal FLAG tag was expressed in *Escherichia coli*, BL21 and purified GS4B affinity, HiTrap SP anion-exchange and Hiload 26/600 S75 size-exclusion chromatographies. The ubiquitylation reaction mixture contained 0.4 µM E1, 6 µM E2, 3 µM E3, 600 µM Ubiquitin, and 100 µM PAF15 in a ubiquitylation reaction buffer [50 mM Tris-HCl (pH 8.0), 50 mM NaCl, 5 mM MgCl_2_, 0.1% Triton X-100, and 2 mM DTT]. The reaction mixture was incubated at 25 °C for overnight.

For *in vitro* deubiquitylation assay, recombinant USP7 full-length wild-type/C223A and deletion of TRAF domain were expressed in Rossetta2 and purified by GST-affinity, HiTrap Q anion-exchange and Hiload 26/600 S200 size-exclusion chromatographies. 3.75 pmol (conc.: 50 nM) of USP7 or USP47 (R&D SYSTEMS, E-626-050) and the ubiquitylated PAF15 were incubated in 75 µl reaction solution in a reaction buffer [20 mM Tris-HCl (pH 7.5), 150 mM NaCl, 0.5 mM DTT and 10% glycerol] at 20 °C for 1 hr. The reaction was stopped at indicated times by adding SDS-sample buffer and the deubiquitylation was analyzed by SDA-PAGE.

### Recombinant proteins expression and purification

GST-PAF15, 3xFLAG-tagged PAF15, 3xFLAG-tagged DNMT1 WT and 4KA mutant, and GST-mDPPA3 61-150 mutant expression and purification were described previously (Mulholland et al., 2020; Nishiyama et al., 2020; Yamaguchi et al., 2017). S79A/S97A, K101A/K105A mutations in pGEX4T-3 and pKS104-PAF15 constructs were introduced using a KOD-Plus Mutagenesis Kit (Toyobo). These mutant x*PAF15* DNAs from pKS104-PAF15 were amplified by PCR and ligated into pVL1392 vector. *USP7 C225S* mutation was also introduced by KOD-Plus Mutagenesis Kit. GST-tagged protein expression in *Escherichia coli* (BL21-CodonPlus) was induced by the addition of 0.1 M Isopropyl-β-D-1-thiogalactopyranoside (IPTG) to media followed by incubation for 12 h at 20 °C. For purification of GST-tagged proteins, cells were collected and resuspended in lysis buffer [20 mM HEPES-KOH (pH 7.6), 0.5 M NaCl, 0.5 mM EDTA, 10% glycerol, 1 mM DTT] supplemented with 0.5% NP-40 and protease inhibitors and were then disrupted by sonication on ice. For FLAG-tagged protein expression in insect cells, 3xFLAG-tagged *USP7 WT* or mutants were transferred from pKS103 vector into pVL1392 vector. Baculoviruses were produced using a BD BaculoGold Transfection Kit and a BestBac Transfection Kit (BD Biosciences), following the manufacturer’s protocol. Proteins were expressed in Sf9 insect cells by infection with viruses expressing 3xFLAG-tagged PAF15 WT or its mutants for 72 h at 27 °C. Sf9 cells from a 750 ml culture were collected and lysed by resuspending them in 30 ml lysis buffer, followed by incubation on ice for 10 min. A soluble fraction was obtained after centrifugation of the lysate at 15,000 xg for 15 min at 4 °C. The soluble fraction was incubated for 4 h at 4 °C with 250 µl of anti-FLAG M2 affinity resin equilibrated with lysis buffer. The beads were collected and washed with 10 ml wash buffer and then with 5 ml of EB [20 mM HEPES-KOH (pH 7.5), 100 mM KCl, 5 mM MgCl_2_] containing 1 mM DTT. Each recombinant protein was eluted twice in 250 µl of EB containing 1 mM DTT and 250 µg/ml 3xFLAG peptide (Sigma-Aldrich). Eluates were pooled and concentrated using a Vivaspin 500 (GE Healthcare).

The human ATAD5-RFC-like complex (ATAD5-RLC) was expressed and purified as follows. Human 293T cells (5 x 10^6^ cells) cultured in a 15 cm dish were transfected with 8 µg of the human mini-AzamiGreen-tagged *ATAD5* gene, and 1 µg each of untagged *RFC2*, *RFC3*, *RFC4*, and *RFC5* genes, all inserted into the pCSII-EF vector, and incubated for 72 h in D-MEM (Sigma-Aldrich) supplemented with 10% fetal bovine serum at 37°C. Cells from ten dishes were harvested, resuspended in 8 ml of PBSGE [140 mM NaCl, 2.7 mM KCl, 10 mM Na_2_HPO_4_, 1.7 mM NaH_2_PO_4_, 20% glycerol, and 20 µM EDTA] supplemented with 1 mM PMSF and 20 µg/ml leupeptin, and lysed by the addition of 0.5% NP-40. The lysates were incubated on ice for 10 min, supplemented with final 0.5 M NaCl, and clarified by centrifugation at 75,000 xg for 30 min at 4 °C. Cleared lysates were then applied onto 1 ml anti-FLAG M2 affinity resin packed in a column equilibrated with PC buffer [50 mM KPO_4_ (pH 7.5), 0.5 mM EDTA, 1 mM 3-[(3-Cholamidopropyl)dimethylammonio]propanesulfonate (CHAPS) and 10% glycerol] supplemented with 0.5 M NaCl. The column was washed with PC buffer supplemented with 0.5 M NaCl, and the ATAD5-RLC proteins were eluted with 100 µg/mL FLAG-peptide (Sigma Aldrich) in the same buffer. Peak fractions were collected and used for the assay.

### Quantification of DNA replication and DNA methylation efficiency in *Xenopus* egg extracts

[α-^32^P] dCTP (3,000 Ci/mmol) and sperm nuclei were added to interphase extracts and incubated at 22 °C. At each time point, extracts were diluted in reaction stop solution (1% SDS, 40 mM EDTA) and treated with Proteinase K (NACALAI TESQUE, Inc.) at 37 °C. The solutions were spotted onto Whatman glass microfiber filters followed by 5% trichloroacetic acid (TCA) containing 2% pyrophosphate. Filters were washed twice in ethanol and dried. The incorporation of radioactivity was counted in the scintillation cocktail. DNA methylation was monitored by the incorporation of S-[methyl-^3^H]-adenosyl-L-methionine. Extracts supplemented with S-[methyl-^3^H]-adenosyl-L-methionine and sperm nuclei were incubated at 22 °C. At each time point, the reaction was stopped by dilution in CPB containing 2% sucrose up to 300 µl. Genomic DNA was purified using a Wizard Genomic DNA Purification Kit (Promega). Incorporation of radioactivity was counted in the scintillation cocktail.

### Immunoprecipitation from chromatin lysate

MNase-digested chromatin fractions were prepared as described previously (Nishiyama et al., 2020). The chromatin pellet was resuspended and digested in 100 µl of digestion buffer [10 mM HEPES-KOH (pH 7.5), 50 mM KCl, 2.5 mM MgCl_2_, 0.1 mM CaCl_2_, 0.1% Triton X-100, 2 mM NEM, 100 µM PR-619] containing 4 U/ml MNase at 22 °C for 20 min. The reaction was stopped by the addition of 10 mM EDTA, and the solution was centrifuged at 17,700 xg for 10 min. For the immunoprecipitation experiment, 2 µg purified IgG, PAF15, USP7 or DNMT1 antibodies were bound to 10 µl of Protein A agarose beads and these beads were mixed with digested chromatin lysates at 4 °C for 2 h. After reaction, these beads were washed by CPB containing 2% sucrose and 0.1% Triton X-100 and resuspended in 10 µl of 2xLaemmli buffer and 20 µl of 1xLaemmli buffer, heated at 100 °C.

## Statistical analysis

### Statistics

The normal distribution of the population at the 0.05 level was calculated using the Shapiro-Wilk normality test. Data are presented as mean ± SEM, unless otherwise noted. Multiple comparisons were performed by Two-way Repeated Measure analysis of variance (RM ANOVA) followed by Sidak’s multiple comparison test. For consistency of comparison, significance in all Figures is indicated as follows. *p < 0.05, **p < 0.01, ***p < 0.001, ****p < 0.0001.

## Data availability

All data generated or analysed during this study are included in the manuscript and supporting files; source data files for all figures have been provided.

## Acknowledgement

We thank Chikahide Masutani for human USP7 antibodies. The study is funded by MEXT/JSPS KAKENHI (JP19H05740 to M.N.; JP19H03143 and JP19H05285 to A.N.; JP16H06578 to M.O.). A.N. was supported in part by a grant from Daiichi Sankyo Foundation of Life Science.

## Supplementary Figures

**Supplemental Figure. S1.**
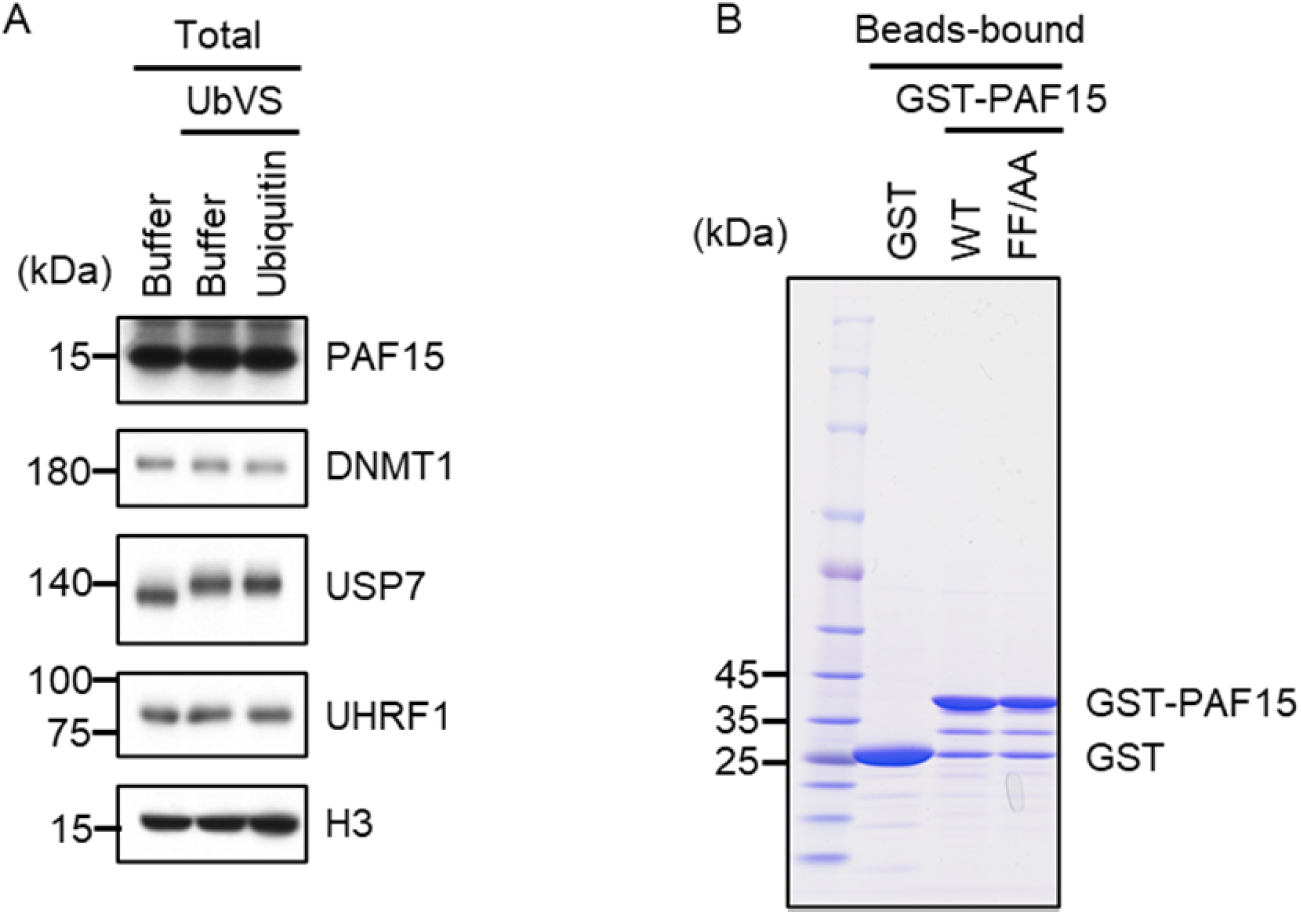
(A) Interphase egg extracts supplemented with UbVS or UbVS+Ub used in Figure 1A were analyzed by immunoblotting using the antibodies indicated. (B) Purified GST or GST-rPAF15 mutants used in Figure 1D were stained using CBB. Source Data are provided as Figure S1-source data.

**Supplemental Figure. S2.**
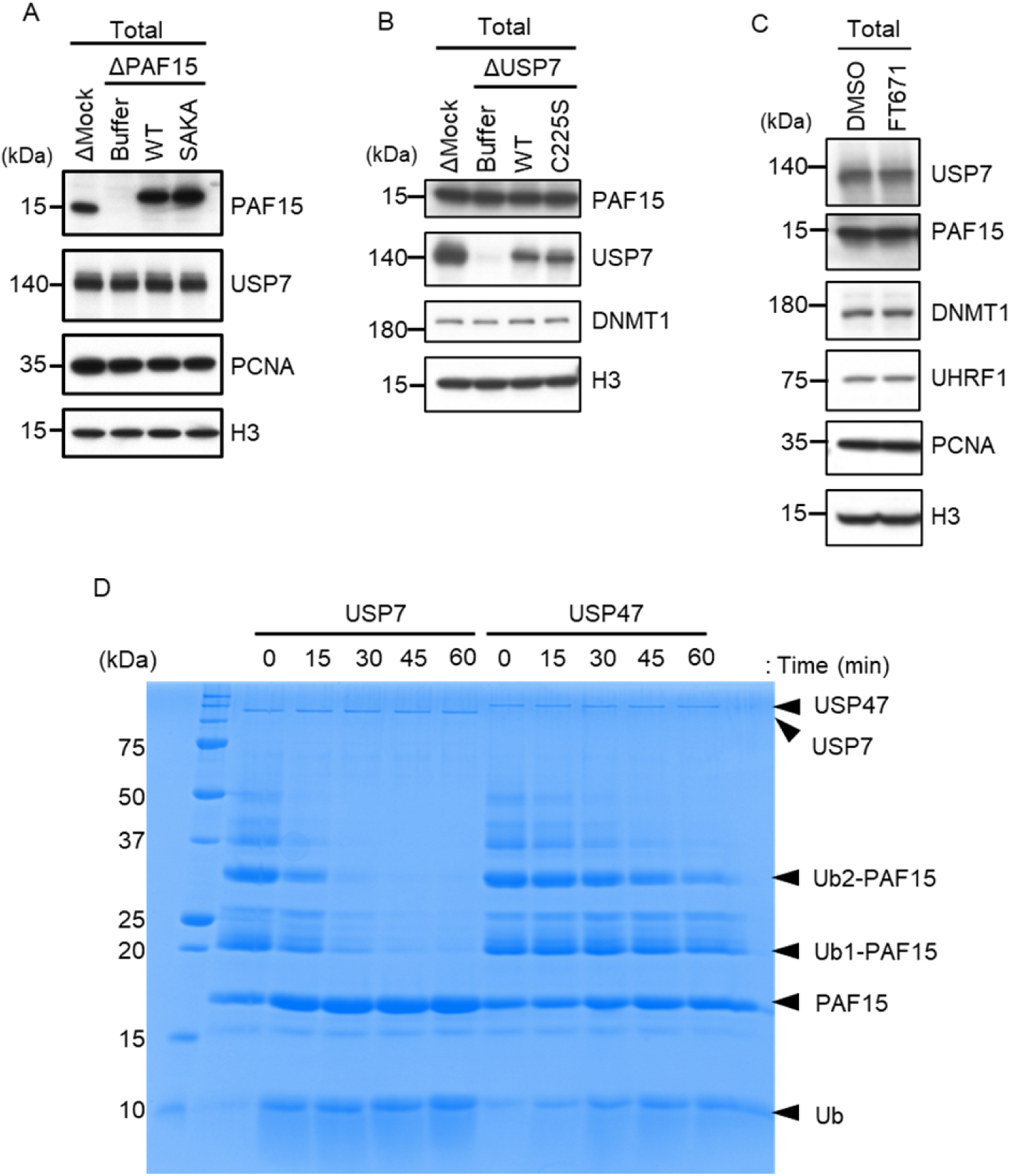
(A) rPAF15 WT-3xFLAG or SAKA-3xFLAG were added to PAF15-depleted extracts. The extracts used in Figure 3A were analyzed by immunoblotting using the antibodies indicated. (B) rUSP7 WT-3xFLAG or C225S-3xFLAG were added to USP7-depleted extracts. The extracts used in Figure 3B were analyzed by immunoblotting using the antibodies indicated. (C) Interphase egg extracts supplemented with Dimethyl sulfoxide (DMSO) or the USP7 inhibitor FT671 used in Figure 3C were analyzed by immunoblotting using the antibodies indicated. (D) Ubiquitinated PAF15 was prepared by *in vitro* ubiquitylation using E1 (mouse UBA1), E2 (UBE2D3), E3 (UHRF1), ubiquitin and C-teminal FLAG tagged PAF15. The reaction mixture was boiled and the precipitant was removed by centrifugation. The supernatant containing ubiquitinated PAF15 was incubated with 50 nM recombinant USP7 or USP47. Incubations were stopped at the indicated time by adding SDS-PAGE loading buffer and analyzed by CBB staining. Source Data are provided as Figure S2-source data.

**Supplemental Figure. S3.**
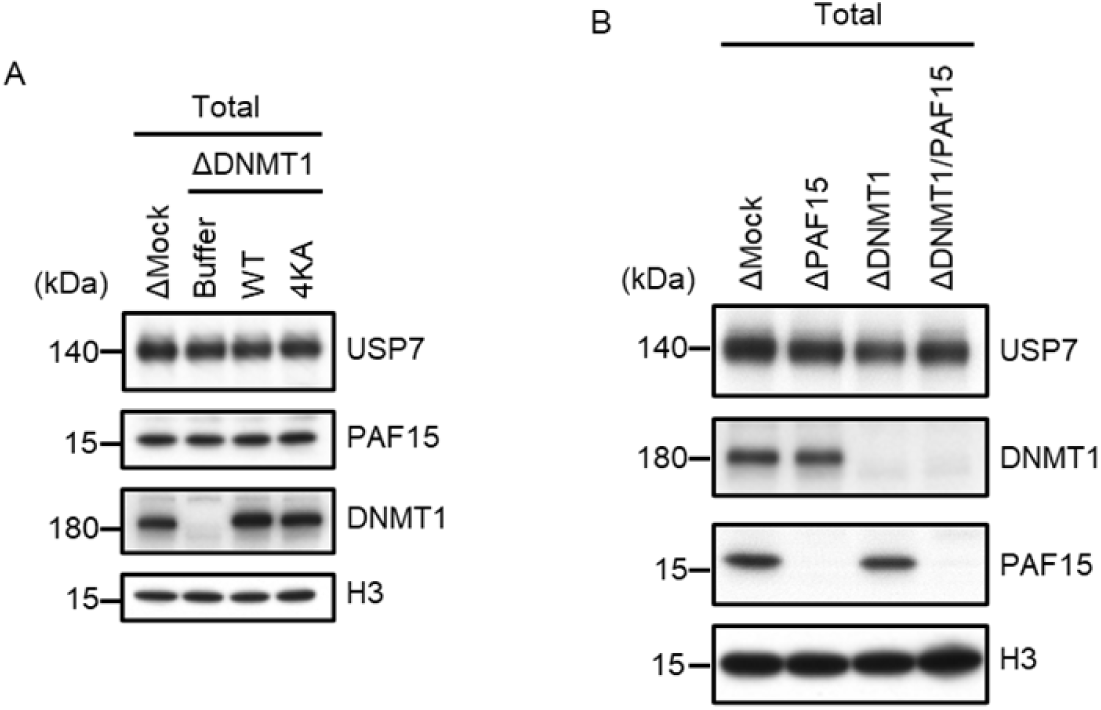
(A) rDNMT1WT-3xFLAG or 4KA-3xFLAG were added to DNMT1-depleted extracts. The extracts used in Figure 4C were analyzed by immunoblotting using the antibodies indicated. (B) Mock-, PAF15-, DNMT1- or PAF15/DNMT1 co-depleted extracts used in Figure 4D were analyzed by immunoblotting using the antibodies indicated. Source Data are provided as Figure S3-source data.

**Supplemental Figure. S4.**
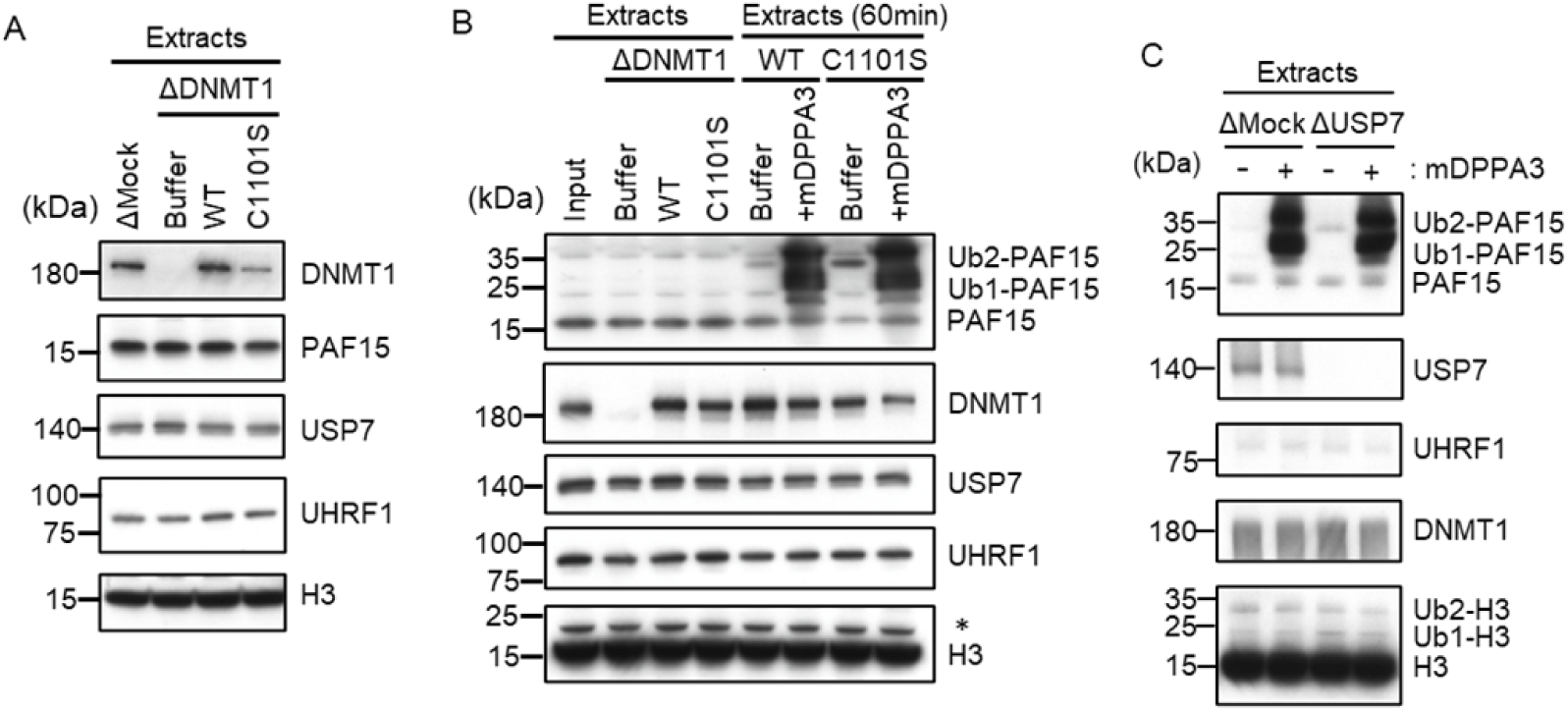
(A) rDNMT1 WT-3xFLAG or C1101S-3xFLAG were added to DNMT1-depleted extracts. The extracts used in Figure 5A were analyzed by immunoblotting using the antibodies indicated. (B) rDNMT1 WT-3xFLAG or C1101S-3xFLAG were added to DNMT1-depleted extracts used in Figure 5D. After 60 min, these extracts were supplemented with GST-mDPPA3. The extracts were analyzed by immunoblotting using the antibodies indicated. The asterisk indicates a non-specific band. (C) Replicating Mock- or USP7-depleted extracts at 90 min were supplemented with either buffer (-) or GST-mDPPA3 (+). The extracts were used in Figure 5E and analyzed by immunoblotting using the antibodies indicated. Source Data are provided as Figure S4-source data.

**Supplemental Figure. S5.**
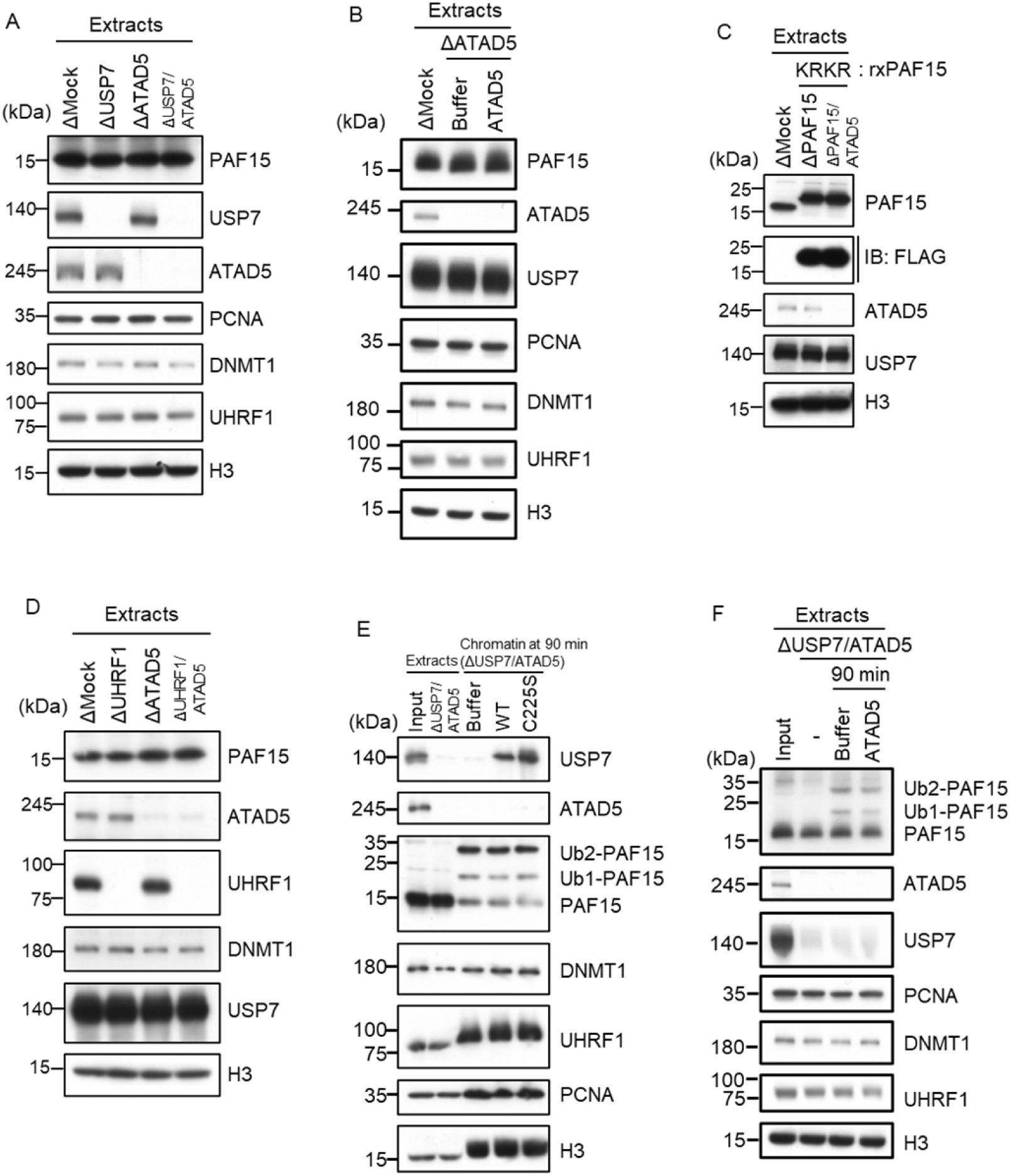
(A) Mock-, USP7-, ATAD5- or USP7/ATAD5 co-depleted extracts were used in Figure 6A and analyzed by immunoblotting using the antibodies indicated. (B) rhATAD5-RFCs was added to ATAD5- depleted extracts. The extracts were used in Figure 6B and analyzed by immunoblotting using the antibodies indicated. (C) rPAF15 KRKR-3xFLAG was added to PAF15- or PAF15/ATAD5 co-depleted extracts. The extracts were used in Figure 6C and analyzed by immunoblotting using the antibodies indicated. (D) Mock-, UHRF1-, ATAD5- or UHRF1/ATAD5 co-depleted extracts were used in Figure 6D and analyzed by immunoblotting using the antibodies indicated. (E)rUSP7WT-3xFLAG or C225S-3xFLAG were added to USP7/ATAD5 co-depleted extracts. The extracts were used in Figure 6E and analyzed by immunoblotting using the antibodies indicated. (F) Replicating USP7/ATAD5 co-depleted extracts at 90 min were supplemented with either buffer or rhATAD5-RFCs. The extracts were used in Figure 6F and analyzed by immunoblotting using the antibodies indicated. Source Data are provided as Figure S5-source data.

**Supplemental Figure. S6.**
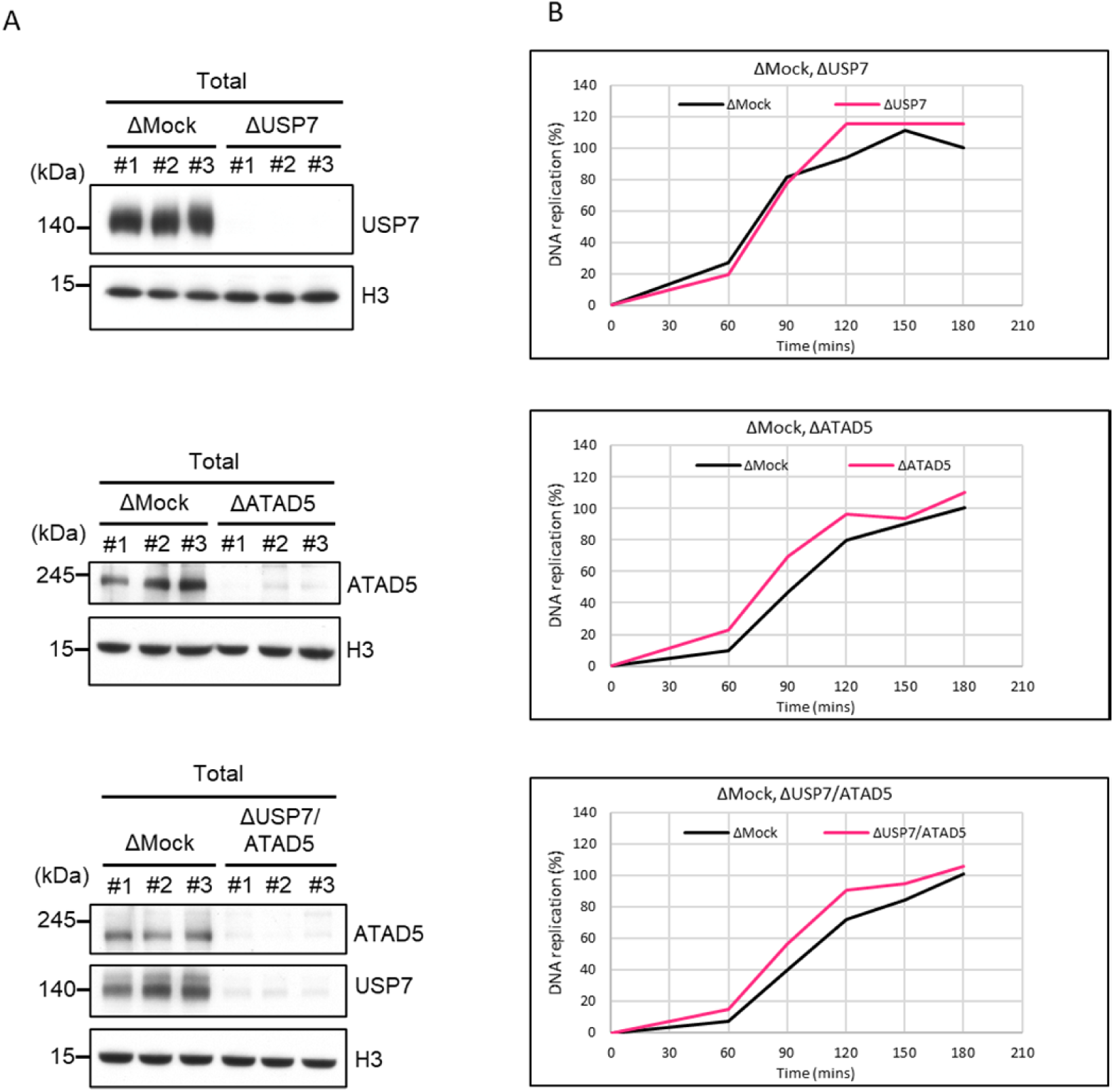
(A) Mock-, USP7-, ATAD5- or USP7/ATAD5 co- depleted extracts used in Figure 7 were analyzed by immunoblotting using the antibodies indicated. (B) Sperm chromatin and radiolabeled [α-^32^P] dCTP were added to Mock-, USP7-, ATAD5- or USP7/ATAD5 co-depleted extracts. Purified DNA samples were analyzed to determine the efficiency of DNA replication. Source Data are provided as Figure S6-source data.

